# Conserved programs and specificities of T cells targeting hematological malignancies

**DOI:** 10.1101/2025.08.07.668919

**Authors:** Tim R. Wagner, Niklas Kehl, Simon Steiger, Tamara Boschert, Michael Kilian, Kane Foster, Gabrielle M. Hernandez, Claudia Ctortecka, Julius Michel, Antonia Schach, Julian Zoller, Anna Metzler, René Onken, Bruno Schönfelder, Simon Renders, Claudia Maldonado Torres, Lilli S. Sester, Jan H. Frenking, Franziska Werner, Wolfram Osen, Katharina Lindner, Ezgi Sen, Sabrina Schumacher, Daria Galas-Filipowicz, Evie Fitzsimons, Edward W. Green, Patrick Schmidt, John M. Lindner, Sebastian Uhrig, Lukas Bunse, Benny Chain, Hartmut Goldschmidt, Niels Weinhold, Stefan Fröhling, Andreas Trumpp, Jennifer G. Abelin, Steven A. Carr, Kwee Yong, Carsten Müller-Tidow, Karsten Rippe, Marc S. Raab, Michael Platten, Stefan B. Eichmüller, Mirco J. Friedrich

## Abstract

T cell-mediated immune surveillance is critical for cancer control, yet its endogenous effectiveness in hematological malignancies remains limited and poorly understood.

Here, we integrate single-cell T cell receptor (TCR) profiling, HLA immunopeptidomics and functional antigen mapping to dissect the specificity landscape of bone marrow lymphocytes (BMLs) in multiple myeloma (MM) and acute myeloid leukemia (AML). We identify a rare subset of tumor-reactive T cells that exhibit a stereotyped transcriptional state distinct from bystander and virus-specific populations. Across both malignancies, immunopeptidomic profiling uncovers a partially conserved antigen repertoire enriched for noncanonical peptides, including products of novel or unannotated open reading frames (nuORFs), pseudogenes, and clonotypic immunoglobulin sequences. Several of these epitopes are recurrently presented and associated with convergent TCR responses across individuals. Based on this immune architecture, we develop a TCR-intrinsic fitness model that infers BML tumor specificity from transcriptional cues and stratifies immunotherapy response across three independent patient cohorts.

Together, these findings map the latent potential of endogenous anti-tumor immunity in two biologically distinct diseases and provide a framework for decoding and restoring productive immune surveillance of hematological malignancies.

**Highlights:**

- Single-cell resolved TCR profiling maps rare tumor-reactive T cells in the bone marrow of multiple myeloma (MM) and acute myeloid leukemia (AML) reveals conserved transcriptional programs
- A shared immunopeptidome across MM and AML includes noncanonical epitopes from nuORFs and idiotype sequences
- Conserved tumor antigens elicit convergent T cell responses across patients

- A TCR fitness model predicts tumor specificity in bone marrow lymphocytes and stratifies immunotherapy response in both hematological malignancies

## Introduction

T cells can mediate durable tumor control, yet their efficacy across hematological malignancies remains highly variable and poorly understood^1–3^. While disease-associated T cells are frequently observed in the bone marrow of patients with multiple myeloma (MM) and acute myeloid leukemia (AML), the frequency, antigen specificity, and clinical relevance of tumor-reactive T cell receptors (TCRs) in these diseases remain elusive^4–6^. Prior studies have demonstrated that most T cells within the tumor microenvironment are bystanders with no overt anti-tumor activity, and that *bona fide* tumor-reactive clonotypes are rare and difficult to identify, particularly outside the setting of immune checkpoint blockade or in the context of highly immunogenic disease^7–9^. In contrast to solid tumors, where the landscape of tumor-specific TCRs and their functional correlates is increasingly defined^10–13^, hematological malignancies lack a unifying model of endogenous antitumor T cell immunity^4–6,14,15^. While neoantigens derived from gene fusions^16,17^ and cancer-associated antigens (CAAs)^18^ have been proposed and tested as actionable targets in selected hematologic malignancies, the prevalence and immunological relevance of shared public antigens or recurrent, cross-disease targets – distinct from private neoantigens – remain incompletely characterized^19–21^.

Here, we integrate high-sensitivity TCR profiling with HLA immunopeptidomes and functional single-cell screening to define the antigen specificity, transcriptional phenotype, and clinical relevance of tumor-reactive bone marrow lymphocytes (BMLs) of patients with MM and AML. By combining antigen-agnostic assays with peptide-guided interrogation of patient-derived T cells, we identify patient-and disease-spanning features of rare tumor-reactive lymphocytes, uncover shared immunogenic peptides derived from both canonical and noncanonical sources, and link the abundance and functional state of these cells to therapeutic response across multiple treatment modalities.

## Results

### Identification of tumor-reactive T cells in the bone marrow of patients with MM and AML

The detection and characterization of tumor-reactive T cells in hematological malignancies is hampered by their rarity and the complexity of the bone marrow microenvironment^22,23^. To identify such clonotypes and define their phenotypic and functional states *in situ*, we performed integrated profiling of bone marrow lymphocytes (BMLs) from a total of 27 patients with newly diagnosed multiple myeloma (NDMM, *n* = 20) and acute myeloid leukemia (AML, *n* = 7). We obtained bone marrow aspirates at initial diagnosis and subjected them to parallel single-cell transcriptomic, proteomic, TCR, and functional analyses (**Figure 1a, Table S1**).

**Figure 1.**
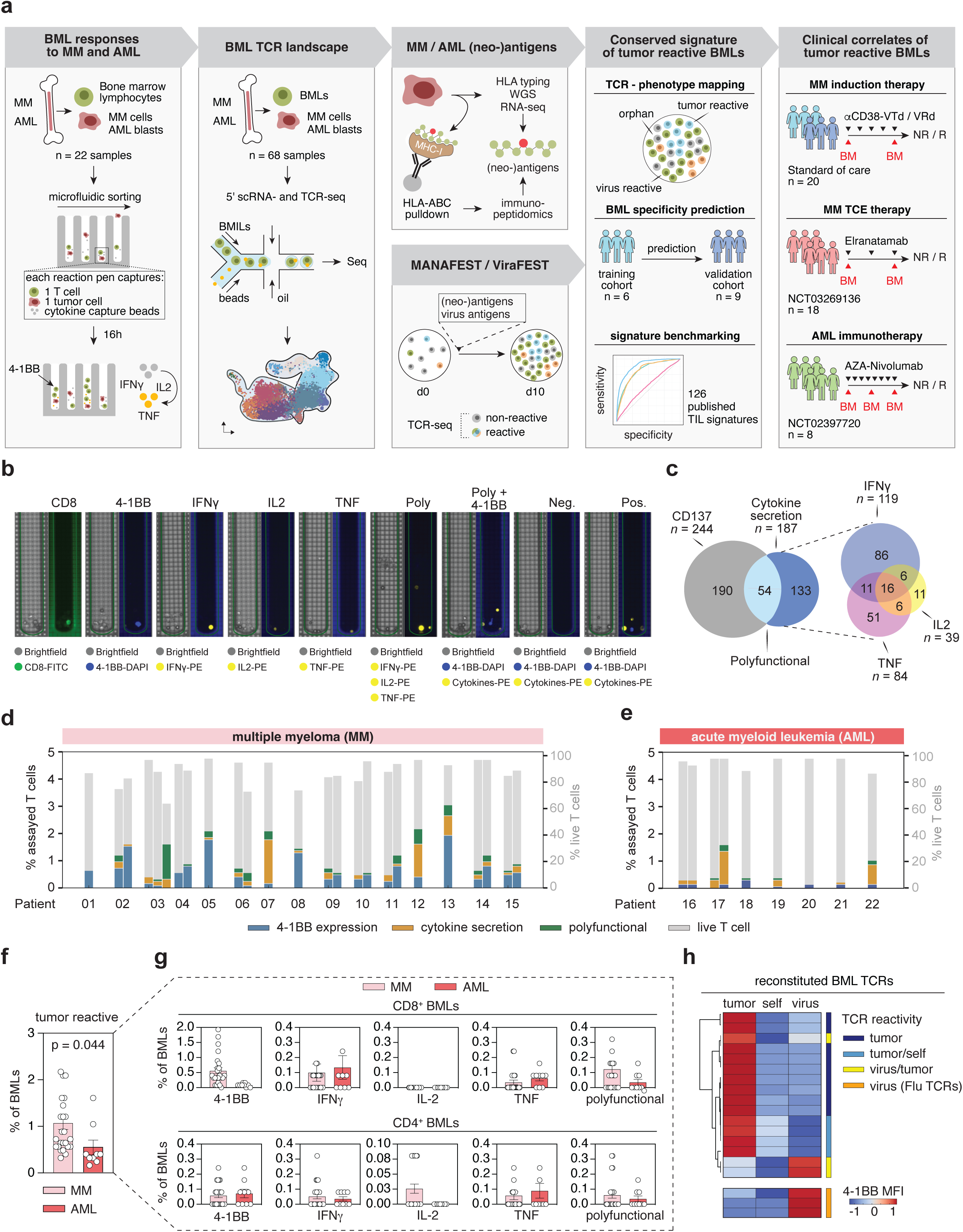
Integrated multi-platform discovery of tumor-reactive T cells and antigens in the bone marrow of patients with multiple myeloma and acute myeloid leukemia **a,** Schematic overview of the pipeline for identifying tumor-reactive bone marrow lymphocytes (BMLs) in multiple myeloma (MM) and acute myeloid leukemia (AML). Steps include: (1) functional co-culture assays with autologous tumor cells in microfluidic chambers, (2) 5′ single-cell RNA and TCR sequencing, (3) tumor antigen discovery via whole genome sequencing (WGS), RNA-seq, and HLA class I immunopeptidomics, (4) Immunopeptidome-informed MANAFEST (mutation-associated neoantigen functional expansion of specific T cells) and ViraFEST assays, (5) TCR-phenotype mapping to single-cell transcriptome data and generation of a conserved transcriptional signature of tumor-reactive BMLs and (6) clinical association with independent patient cohorts and therapeutic settings. **b,** Representative images from the functional screening assay. Single BMLs were co-cultured with autologous tumor cells (MM / AML) for 16 hours in nanowell chambers containing cytokine-capture beads and monitored for 4-1BB, IFN-γ, IL-2, and TNF expression. Tumor-reactive cells were retrieved and subjected to TCRA/B sequencing. **c,** Venn diagram summarizing characteristics of 673 tumor-reactive TCRs. Polyfunctionality was defined as concurrent expression of 4-1BB and ≥1 cytokine. **d-e,** Distribution of tumor-reactive events across 25 MM (d) and 9 AML (e) assay runs. Functional categories defined as in (c). **f,** Cumulative frequency of tumor-reactive BMLs per sample. Statistical comparison between MM and AML was performed using an unpaired two-tailed t-test. **g,** Proportion of tumor-reactive BMLs stratified by CD4⁺ or CD8⁺ T cell subsets. Statistical comparison between MM and AML was performed using unpaired two-tailed t-tests. **h,** Heatmap summarizing tumor specificity testing for 19 representative reconstituted BML-derived TCRs, each tested against autologous tumor cells (tumor), healthy PBMCs (self), and a viral peptide pool (virus). Data represent mean 4-1BB MFI across n=3 experimental replicates per TCR.

To establish a high-resolution map of BML phenotypes and their TCR clonotypes, we applied single-cell RNA sequencing (scRNA-seq) combined with single-cell V(D)J sequencing (scVDJ-seq), enabling us to simultaneously assess gene expression and paired TCR identity (**Figure S1a-d**). We recovered 187,015 high-quality transcriptomes representing 132,501 unique TCR clonotypes across the patient cohort (**Figure S2a-d**). Unsupervised clustering of BMLs revealed 13 phenotypic subsets, including major populations of CD8⁺ effector-memory (CD8_EM), progenitor-exhausted (CD8_PEX), and stem-like memory (CD8_SM) T cells, as well as naïve and memory CD4⁺ subsets and regulatory T cells, consistent with previous studies in untreated MM and AML^24–28^ (**Figure S1c-d, Figure S2a-b, Table S2**). Minor populations included NK-like and quiescent T cells.

To link phenotype with clonal architecture, we analyzed 132,501 TCR clonotypes at initial diagnosis. Expanded clones were enriched with cells in CD8_EM and CD8_PEX phenotypes, consistent with antigen-driven selection (**Figure S2c-f**). In contrast, clonally diverse and non-expanded T cells primarily occupied naïve and memory CD4⁺ compartments (**Figure S2g**), suggesting that clonal expansion alone associates with the state and lineage of BMLs but may not be a sufficient indicator of tumor reactivity.

To identify functionally tumor-reactive T cells in both diseases, we used a dual screening strategy: 1) A forward screen exposing single BMLs to patient-autologous MM/AML cells in microfluidic chambers; and 2) A reverse antigen-informed screen based on WGS/RNA-sequencing-predicted and immunopeptidomics-defined tumor antigens, applying a frequently used functional expansion assay (MANAFEST/ViraFEST) to detect antigen-responsive T cells^10,29–31^ (**Figure 1a**).

In the forward screen, single BMLs were co-incubated with autologous primary tumor cells for 16 hours. The microfluidic assay simultaneously captured cytokine secretion (IL-2, IFN-γ, TNF) and surface upregulation of CD137 (4-1BB) upon tumor cell contact (**Figure 1b**). Given the median count and distribution of unique BMLs clonotypes in our patient biopsies by the concurrently performed scTCR-seq and considering the benchmarks of our screening approach, the minimal frequency of an individual TCR to be detected at least once with ≥ 95% probability is approximately 0.2% (or 42 cells per clone). Across 42,262 BMLs functionally screened, we identified 673 BMLs (∼1.21% per patient) demonstrating at least one signal of tumor reactivity, which were retrieved and subjected to paired TCRA/B sequencing (**Figure 1b-c, Table S3, Supplementary Data File 1**). Interferon gamma (IFN-γ) was the most frequently secreted cytokine, followed by Tumor Necrosis Factor (TNF) and Interleukin-2 (IL-2). A subset of T cells exhibited polyfunctionality, with simultaneous cytokine secretion and CD137 expression (**Fig. 1c**). Tumor-reactive BMLs were detected in nearly all patients, but their abundance and functional profiles varied across individuals and disease entities (**Figure 1d-e, Table S3**). MM samples displayed a higher overall frequency of tumor-reactive cells than AML, particularly among CD8⁺ polyfunctional T cells co-expressing CD137 (**Figure 1f-g, Table S3**). IL-2 only-secreting cells were exclusively CD4⁺, and we were able to retrieve and identify CD4^+^ as well as CD8^+^ TCRs with a progenitor-exhausted phenotype (**Figure 1g, Figure S3a**). Using the CDR3-sequence as a unique clonal barcode to map these functionally identified TCRs to the scRNA/V(D)J-seq dataset^32,33^, we found that the abundance of tumor-reactive clones correlated with canonical transcriptional programs. Less expanded tumor-reactive T cells expressed lymphoid progenitor markers such as *SELL* and *IL7R*, while highly expanded reactive clones expressed cytotoxic NK-like effector genes including *GZMB*, *GNLY*, *PRF1*, and *NKG7* (**Figure S3b**).

To validate the tumor specificity of recovered TCRs, we reconstructed selected clonotypes via mRNA-based TCR re-expression in patient-autologous peripheral blood T cells (**Figure S3c-g**). A subsequently performed orthogonal co-culture assay used upregulation of CD69 to validate HLA-dependent TCR responses in each patient against autologous tumor cells but not against autologous healthy PBMCs or viral peptides, though a subset of TCRs displayed cross-reactivity (**Figure 1h, Figure S4a-d, Figure S5a-b, Figure S6a-c**). Taken together, these functional screening data suggest the prevalence of tumor-reactive TCRs among BMLs in both MM and AML.

### Noncanonical immunopeptidomes and shared antigen vulnerabilities in MM and AML

Having identified tumor-reactive TCRs with selective activity against autologous malignant cells, we next aimed to delineate the antigenic landscape that underlies these specificities *in vivo*. Building on emerging insights into the immunogenic potential of aberrant transcript processing^34,35^, including recent evidence of mis-splicing-derived neoepitopes in splicing factor-mutant leukemias, we hypothesized that hematological malignancies, despite their comparatively low tumor mutational burden relative to solid tumors, may give rise to immunogenic, noncanonical antigens as a byproduct of transcriptional and translational dysregulation^21^. To explore this, we conducted systematic HLA class I immunopeptidomics on primary tumor samples from 11 patients with MM and 6 patients with AML in our cohort, each informed by prior HLA typing (**Table S4**). Using LC-MS/MS analysis of HLA-I immunoprecipitates, we identified 17,893 unique peptide sequences across the cohort. Integration with curated annotations of upstream/downstream open reading frames (ORFs), long noncoding RNAs (lncRNAs), pseudogenes, and cancer-associated antigens (CAAs) revealed that a substantial non-healthy proportion of HLA-I-bound peptides were indeed derived from noncanonical translational events^36^ (**Figure 2a, Tables S5-6**). These peptides were observed across all samples and accounted for a major fraction of the tumor immunopeptidome. The number and composition of presented peptides varied markedly between patients and entities, underscoring the high inter-individual heterogeneity of the HLA-I presented immunopeptidome (**Figure 2a**). Although CAAs were identified in both diseases, they contributed a modest fraction of the total peptide pool (∼5-10%) and were more frequently detected in MM^37^ (**Figure 2a, Table S6)**.

**Figure 2.**
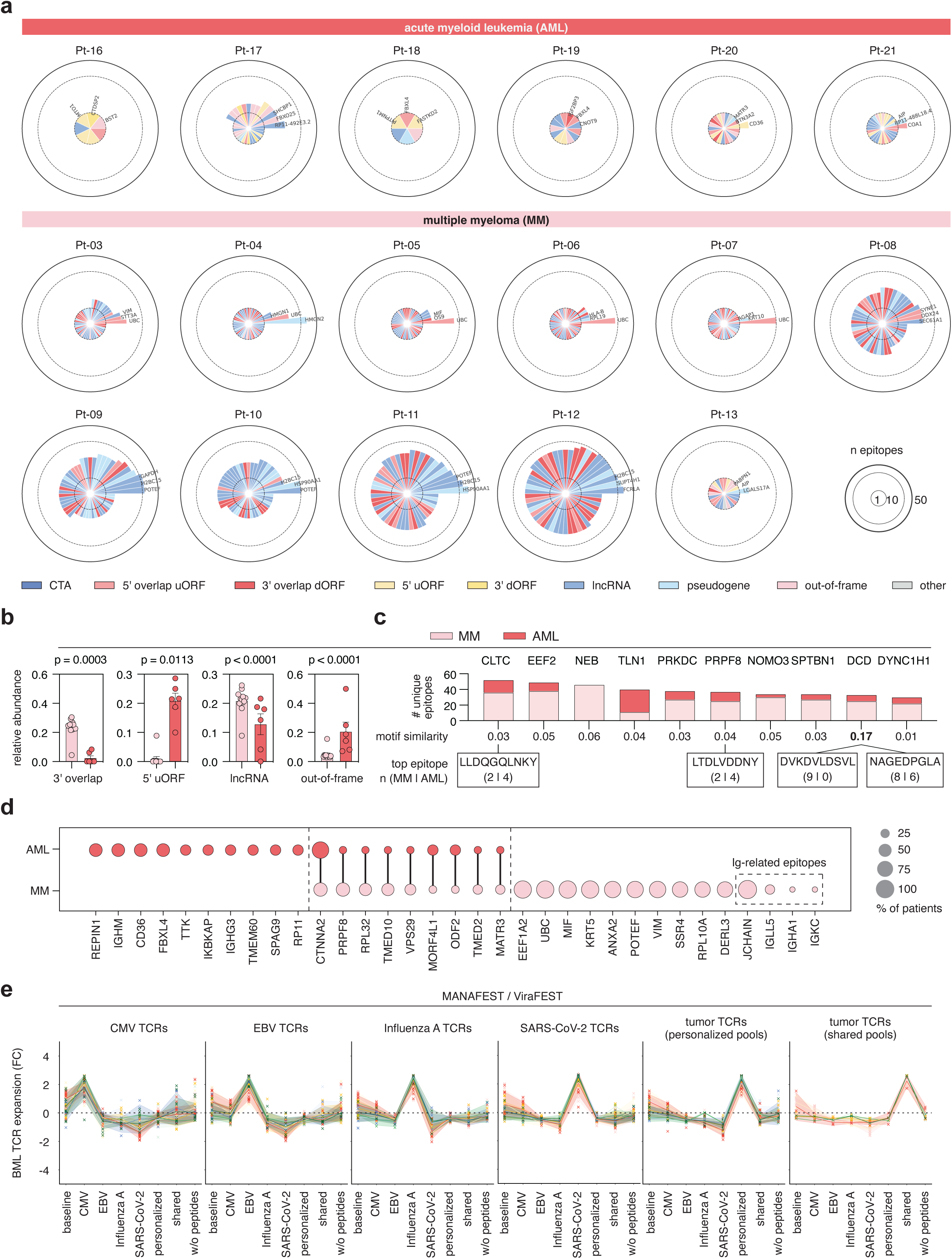
Non-canonical antigen landscapes in MM and AML and cross-entity epitopes revealed by HLA immunopeptidomics **a,** Circular bar plots illustrating the diversity and source of non-canonical HLA class I peptides identified in individual AML (top) and MM (bottom) patients. Epitope sources include cancer-testis antigens, upstream/downstream non-canonical ORFs, lncRNAs, pseudogenes, and others. **b,** Bar plots comparing the relative abundance of selected non-canonical epitope categories in MM vs. AML. Statistical comparisons performed using unpaired two-tailed t-tests. **c,** Bar plots showing the number of unique epitopes derived from abundant non-canonical source proteins across MM and AML patients. **d,** Bubble plot summarizing the frequency of epitope source proteins across patients and disease entities; circle size reflects the percentage of patients with detected epitopes from each protein. **e,** Summary TCR expansion data of MANAFEST (mutation-associated neoantigen functional expansion of specific T cells) and ViraFEST assays across 9 patients with matched immunopeptidomes. Only significantly enriched TCRs per condition, identified by bootstrapped enrichment (FEST algorithm), are shown.

The noncanonical antigen landscape diverged systematically between MM and AML. AML samples predominantly presented peptides from 5′ untranslated region-derived uORFs, while MM samples were enriched for 3′ overlapping dORFs, lncRNA-encoded peptides, and pseudogene-derived sequences. These differences remained statistically significant after normalization for total peptide counts (**Figure 2b, Table S5**).

However, we identified a subset of noncanonical peptides that were recurrently detected across both diseases. For example, a distinct peptide from *DCD*, a gene expressed in normal epithelial tissues and aberrantly activated in hematological cancers, was HLA-presented in 8 of 11 MM patients and all 6 AML samples^38^. Additional peptides from *CTNNA2*, *ODF2*, and members of the zinc finger (ZNF) family were shared across disease types, echoing recent reports of conserved HLA-presented peptides as a result of splicing factor-mutations^21,34,35^ and suggesting that convergent antigen processing pathways may arise across genetically and clinically distinct malignancies, potentially through shared stress or epigenetic mechanisms that deregulate RNA translation (**Figure 2c-d**).

Beyond nuORF-derived peptides, we also identified several peptides originating from the immunoglobulin protein complex in MM samples as HLA-I-bound antigens presented by malignant plasma cells (**Figure 2d**). To systematically investigate this phenomenon in MM, we performed *in silico* assembly of the malignant B cell receptor (BCR) repertoire by reconstructing the dominant myeloma clonotype in each patient. We then predicted peptide sequences with high binding affinity to each patient’s autologous HLA class I allotypes and generated individualized BCR-derived peptide libraries for targeted immunopeptidomic mapping (**Figure S7a, Table S7**). In one patient (Pt-11), we thereby identified a naturally presented peptide (LYKFNNKEYL) derived from the hypermutated *IGKV* chain, specifically within the complementarity-determining region 1 (CDR1). This peptide was eluted from the HLA class I complex and matched a variant detected at 40% variant allele frequency in the dominant MM clone of the patient, providing direct evidence of BCR-derived peptide presentation in MM (**Figure S7b**). Together with prior reports on immunoglobulin-derived antigens in lymphomas and idiotype vaccinations, these data underscore the potential of clonotypic immunoglobulin sequences as an alternative personalized neoantigen source uniquely accessible in B cell-derived malignancies^39–45^.

To complement our forward TCR screening approach and broaden coverage of the BML specificity landscape, we next sought to retrospectively identify TCRs reactive to the identified tumor antigens as well as those recognizing common viral epitopes. We performed MANAFEST and ViraFEST assays in 9 patients for whom both immunopeptidomic data and viable BMLs were available^10,29–31^. T cells were stimulated with personalized peptide pools comprising immunopeptidomics derived putative private and shared tumor antigens, alongside defined viral epitope libraries. Following 10-day culture and ultradeep TCRβ sequencing, we identified 123 tumor antigen-specific (MANA-specific) TCRs across all individuals: 88 reactive to private antigens and 35 targeting shared tumor epitopes. In parallel, we detected 104 CMV-specific, 86 EBV-specific, 88 Influenza A-specific, and 88 SARS-CoV-2-specific TCRs (**Figure 2e, Table S8**). No overlap was observed between tumor-reactive and virus-specific TCRs. Convergent TCR selection is well established in the context of viral immunity, where population-wide exposure to conserved epitopes drives the emergence of public TCRs^46,47^. In addition to virus-specific TCRβ sequences expanding similarly across individuals in response to common viral epitopes, we also identified several shared TCRβ sequences across patients in response to common tumor-derived peptides in the MANAFEST assay, suggesting that a similar phenomenon may occur in response to recurrently presented tumor antigens (**Table S8**).

Together, these findings reveal a complex yet structured tumor antigen landscape in AML and MM, shaped by disease-specific translational programs and partially conserved epitopes. The recurrent presentation of nuORF-derived peptides and the natural display of clonotypic, idiotype-derived neoepitopes suggest alternative sources of tumor-specific peptides for TCR recognition in the absence of frequent mutation-derived neoantigens.

### Mapping of tumor-specific TCRs and convergent selection

To resolve antigen specificities of the tumor-reactive TCRs identified in our screens, we next performed a peptide library-based TCR:epitope deconvolution approach (adapted from the T-FINDER system^48^). We assembled personalized antigen libraries composed of patient-specific and shared peptides identified by immunopeptidomics, additionally including predicted BCR-derived neoepitopes in MM cases. Lentivirally transduced Jurkat reporter cells expressing individual tumor-reactive TCRs were co-cultured with patient-derived B lymphoblastoid cell lines (B-LCLs), which served as HLA-matched autologous antigen-presenting cells (**Figure S8a**). Out of 32 assembled tumor-reactive TCRs, 31 were successfully reconstituted in the reporter platform, with one clone excluded due to unstable expression (**Figure S8b-c**). Using CD69 upregulation as a readout of TCR activation, and comparing responses to irrelevant peptide controls, we successfully deorphanized 21 TCRs with confirmed antigen specificities (**Figure S8d-f, Table S9**). We further investigated endogenous T cell recognition of the *IGKV*-derived neoepitope identified in Pt-11 using this approach and recovered three independent endogenous TCR clonotypes capable of recognizing different minimal epitopes of the naturally presented *IGKV* peptide (LYKFNNKEYL) (**Figure S7c, Tables S7, S9**).

To map tumor-specific TCRs within the broader immune architecture of the bone marrow, we next integrated the functional data of our antigen-agnostic and antigen-informed assays with baseline BML transcriptomes. By leveraging the unique CDR3 nucleotide sequences of validated TCRs as clonal barcodes, we retrospectively traced them to their baseline transcriptional states in our scRNA/V(D)J-seq dataset of BMLs. This approach enabled us to resolve the native phenotypes of tumor-reactive T cells independent of stimulation-induced transcriptional perturbation^32,33^ (**Figure 3a, Figure S2**).

**Figure 3.**
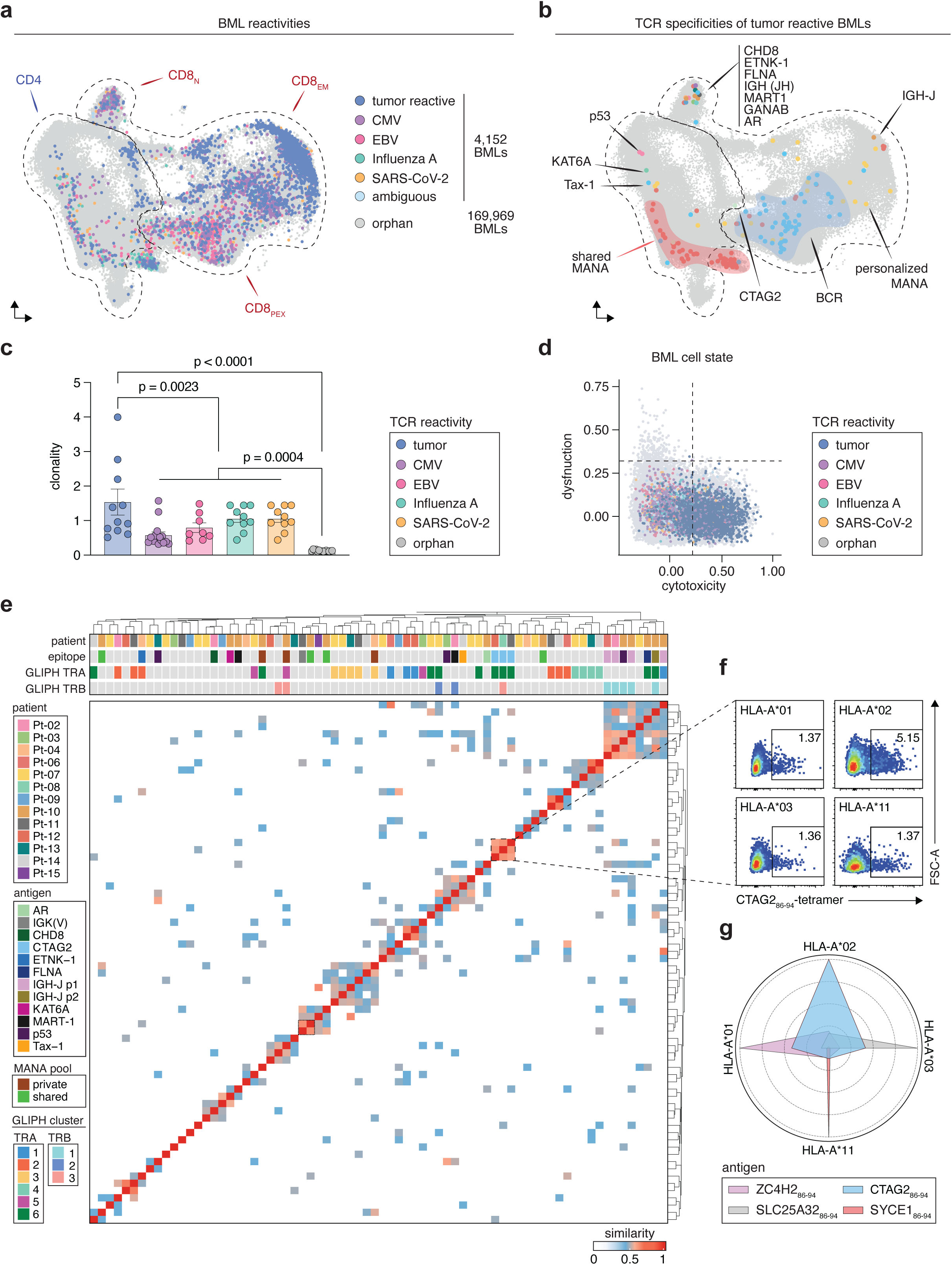
Mapping antigen specificity and diversity across the bone marrow T cell landscape **a,** UMAP of BMLs colored by TCR specificity: tumor-reactive (deorphanized, MANA-specific, or antigen-agnostic), virus-specific (SARS-CoV-2, Influenza A, CMV, EBV), ambiguous (virus/tumor or multiple viruses). Orphan cells in grey. **b,** UMAP of deorphanized BMLs colored by matched peptide antigen (see Figure S7-9). **c,** TCR clonality (1/Shannon diversity) stratified by TCR specificity group. Statistical comparisons performed using one-way ANOVA with Tukey’s post hoc test. **d,** Scatter plot of cytotoxicity versus dysfunction module expression across BMLs, color-coded by antigen specificity. Bystanders in gray. **e,** Heatmap of pairwise TCR CDR3αβ sequence similarity using BLOSUM45-or GLIPH-based scoring. Only values exceeding the 95th percentile bootstrap threshold are shown; rows annotated by patient origin. **f,** Flow cytometry gating for peptide-MHC tetramer staining using CTAG2₈₆-₉₄ (RLLELHITM) presented on representative HLA-A alleles (*01, *02, *03, *11). **g,** Radar plot showing binding promiscuity of shared immunopeptidome-derived antigens across HLA-A supergroups, based on empirical tetramer reactivity.

Across 15 patients, we back-projected 2,249 T cells corresponding to 100 tumor-reactive clonotypes. Of these, 46 TCRs were linked to defined peptide antigens with a corresponding high-quality transcriptome (**Figure 3b, Table S10**). Tumor-reactive T cells within the single cell sequencing dataset constituted approximately 1.29% of all bone marrow-infiltrating T cells at diagnosis, aligning with frequencies observed in our functional screening as well as in solid tumors^7,11^. In contrast, the majority of TCRs not classified as tumor-reactive mapped to CD4⁺ T cells or matched empirically defined viral antigen-recognizing TCRs (CMV, EBV, Influenza A, and SARS-CoV-2). These distributions are consistent with published descriptions of virus-specific immune landscapes in healthy bone marrow^49–51^ (**Figure 3a-b**). Tumor-specific TCRs exhibited significantly higher clonality than viral-or orphan TCRs (**Figure 3c**), suggesting antigen-driven local expansion. These expanded clones also displayed transcriptional states associated with cytotoxicity (**Figure 3d**). In contrast, virus-specific TCRs largely resembled non-expanded bystanders with lower cytotoxic potential and a more quiescent phenotype, as previously reported^7,52^ (**Figure 3c-d**).

Given that certain tumor antigens were recurrent across patients, we next investigated whether chronic exposure to shared tumor-associated epitopes could lead to convergent selection - the emergence of TCRs with similar antigen-binding motifs across individuals^46,47^. We defined convergent TCRs as those that shared at least one V gene and a homologous CDR3 amino acid sequence with another patient-derived clone, irrespective of nucleotide sequence (i.e., “public” rearrangements). After applying bootstrapped similarity thresholds and motif clustering and cross-representation of our approach with GLIPH^53^, we identified multiple TCR networks exhibiting high inter-patient similarity, indicative of convergent recognition patterns (**Figure 3e, Figure S9a**).

Among the tumor-reactive TCRs, we identified two representative clusters of convergent clonotypes. The first comprised three multiple myeloma patients (Pt-02, Pt-06, Pt-12) sharing a homologous TCRβ CDR3 sequence specific for the immunoglobulin heavy-chain joining segment (*IGHJ*), a tumor-associated antigen highly expressed in plasma cell malignancies but absent in healthy non-B cells^54^ (**Figure 3e**). The second cluster featured a shared TCRα CDR3 motif across Pt-06, Pt-08, and Pt-11, with variable yet overlapping TCRβ sequences (**Figure 3e, Figure S9a**). Using HLA profiling data and tumor cells from patient Pt-08, we validated *CTAG2*₈₆₋₉₄ (RLLELHITM) as the cognate antigen of this clonotype, confirming antigen specificity across patients with non-overlapping HLA haplotypes (**Figure S9b-c**). To assess the breadth of this shared reactivity, we performed tetramer staining for *CTAG2*₈₆₋₉₄ and three other recurrent HLA class I-presented epitopes using BMLs from patients with four distinct HLA haplotypes (**Figure S10a-b**). *CTAG2*₈₆₋₉₄-reactive T cells were detected in multiple individuals across different HLA supergroups, with a preferential but not exclusive binding to HLA-A*02 (**Figure 3f, Figure S10b**). These data indicate HLA promiscuity of the *CTAG2*₈₆₋₉₄ epitope, a property of some antigens that allows cross-patient immune recognition despite HLA diversity^55^. Notably, the frequency of *CTAG2*₈₆₋₉₄-specific T cells exceeded that of other shared antigens tested, suggesting its potential as an actionable target (**Figure 3f-g**).

Together, these data suggest that tumor-specific T cell responses in MM and AML are not entirely private but can involve public TCR motifs and shared tumor antigens with broad HLA-restricted epitope presentation.

### Conserved transcriptional programs define tumor-reactive T cells in the bone marrow

To determine whether tumor-reactive BMLs share conserved transcriptional features that distinguish them from virus-specific or bystander T cells, we performed differential gene expression analysis between tumor-reactive, virus-reactive and orphan BMLs. Tumor-reactive T cells displayed a predominant effector-memory phenotype, marked by expression of cytotoxic effector molecules (*GZMB*, *GZMH*, *GNLY*) and tissue-associated markers (*ZNF683*, *CCL4*), consistent with a microenvironment shaped by chronic antigen stimulation. These features were not shared with virus-reactive TCRs. Notably, CMV-and EBV-reactive cells downregulated transcriptional regulators *ZBTB21*^56^ and *USP22*^57^, while influenza-specific T cells showed enrichment for proliferation-and stress-related genes but lacked overt features of cytotoxicity. By contrast, tumor-reactive clones uniquely co-expressed effector molecules with exhaustion markers (**Figure 4a-b**).

**Figure 4.**
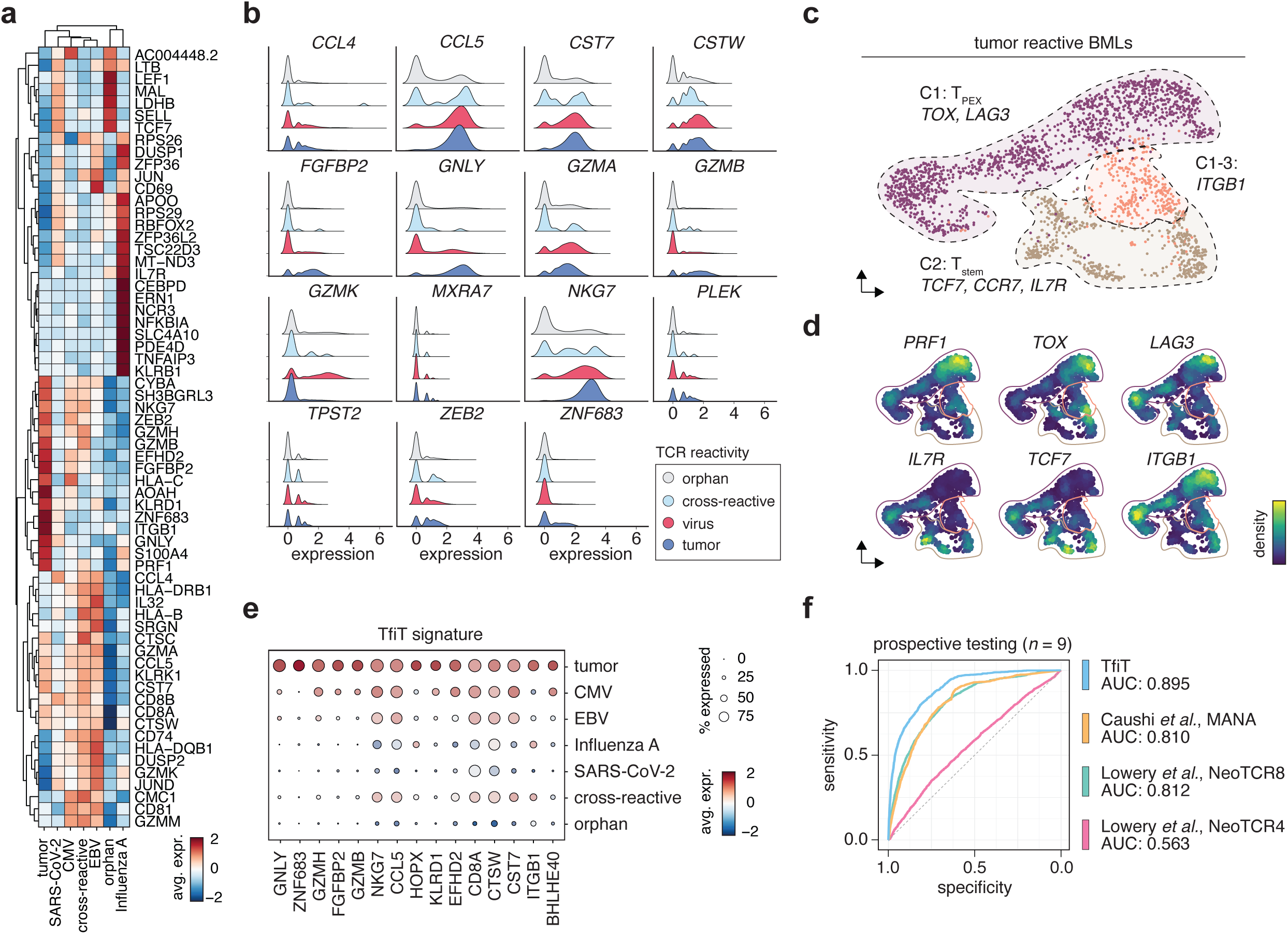
A conserved transcriptional signature of tumor-reactive bone marrow T cells **a,** Scaled heatmap of top differentially expressed genes per T cell reactivity category across 15 patients. **b,** Ridge plots of representative marker genes stratified by reactivity category: tumor, virus, ambiguous (tumor/virus), and bystander. **c,** UMAP of 2,249 validated tumor-reactive cells representing 100 unique TCR clonotypes. **d,** UMAPs overlaid with expression density for selected marker genes. **e,** Dot plot of TfiT signature gene expression across reactivity categories in the signature discovery cohort (n=6 patients). **f,** AUROC analysis comparing predictive performance of transcriptional classifiers for the identification of tumor-reactive TCRs: TfiT (AUC=0.895), MANA_Caushi^10^ (0.810), NeoTCR_8^12^ (0.812), and NeoTCR_4^12^ (0.563). Predictions evaluated in a prospective cohort (n=9 patients; 101,256 cells; 1,302 functionally validated tumor-reactive BMLs).

To further dissect this transcriptional phenotype of tumor-reactive BMLs, we performed re-clustering of TCRs with experimentally determined tumor reactivity only, which revealed three dominant transcriptional states (**Figure 4c-d**): 1) TR-TPEX: a progenitor-exhausted-like phenotype co-expressing *TOX* with cytotoxic mediators (*GZMB*, *PRF1*), reminiscent of CD8⁺ T cells with long-term persistence and therapeutic relevance in chronic infections and cancer; 2) TR-TSTEM: a stem-like memory population characterized by high expression of *TCF7*, *CCR7*, and *IL7R*, and low expression of exhaustion or effector genes, suggestive of a less-differentiated, renewal-capable state; 3) An intermediate TR cluster representing transitional states between these two axes. While individual tumor-reactive clonotypes could span multiple phenotypes, the majority (61.8%) of cells clustered within the TR-TPEX state, reflecting the dominant differentiation trajectory of antitumor CD8⁺ T cells within the bone marrow. A smaller fraction (23.5%) resided in the TR-TSTEM state, suggesting a small reservoir of these stem-like tumor-specific precursors at initial diagnosis. In line with this distribution, the most clonally expanded tumor-reactive TCRs - those comprising >1% of T cells in baseline bone marrow biopsies - were predominantly found within TR-TPEX and CD8_EM compartments (**Figure S3b**), consistent with antigen-driven differentiation and localized expansion.

To systematically define the transcriptional features of tumor-reactive bone marrow lymphocytes (BMLs), we generated a classifier based on genes that consistently distinguished tumor-specific from virus-specific T cells in our patients. Using iterative feature selection, model training, and retrospective ROC analysis on a reference dataset of 947 tumor-reactive cells from six patients, we identified a robust 15-gene signature with high predictive accuracy (AUC = 0.979) (**Figure 4e, Table S11**). We termed this classifier TFiT (Tumor-reactive Features in T cells). When benchmarked on the same dataset, TFiT outperformed 122 existing signatures derived from neoantigen-or mutation-associated neoantigen (MANA)-specific T cells, including MANA^10^ (Caushi et al., AUC = 0.810), NeoTCR_8^12^ (Lowery et al., AUC = 0.812), and NeoTCR_4^12^ (Lowery et al., AUC = 0.563) (**Figure S11a**^10–13,58–61^**, Tables S11-12**). Notably, whereas prior signatures were primarily trained on tumor-infiltrating lymphocytes (TILs) from solid tumors, TFiT is derived from BMLs, which might reflect the unique set of microenvironmental cues in the bone marrow niche and therefore underlie its superior performance in this context^62,63^.

To evaluate its prospective performance, we applied TFiT to an independent validation cohort of 1,302 T cells from nine additional patients. TFiT again outperformed all comparator models, achieving an AUC of 0.895 (**Figure 5f, Figure S11b, Table S12**), confirming its generalizability across patients and disease entities in identifying tumor reactive BMLs.

**Figure 5.**
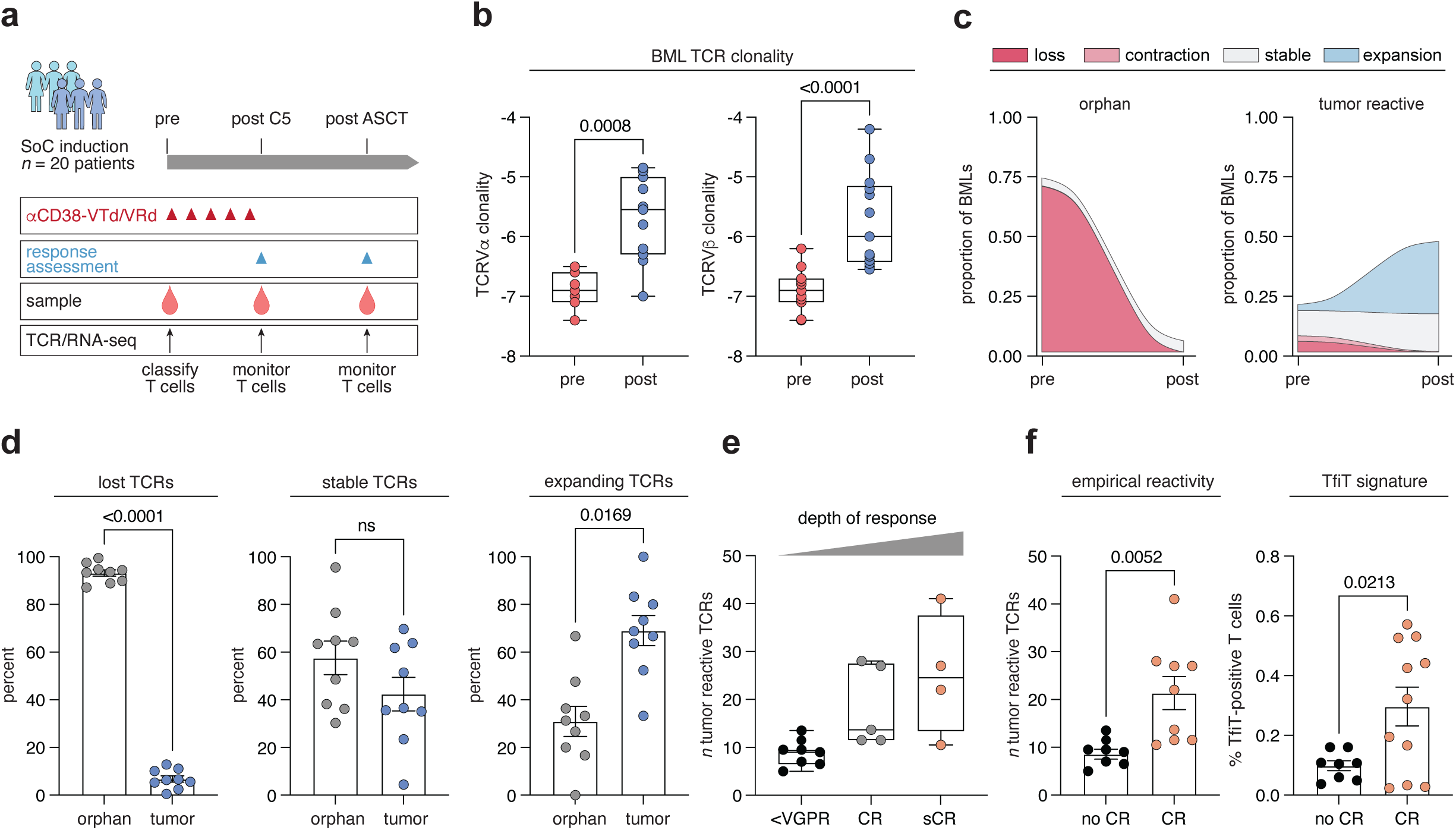
Longitudinal dynamics and clinical relevance of tumor-reactive BMLs during MM induction therapy. **a,** Experimental workflow and sampling timeline for 20 newly diagnosed MM (NDMM) patients treated with Dara-VTd followed by ASCT. BMLs were profiled by scRNA-seq and scTCR-seq pre-treatment, post-induction, and post-ASCT. **b,** TCR clonality (1/Shannon diversity) based on TRAV/TRBV usage, assessed pre-treatment and 100 days post-ASCT from the CARDAMON-Trial^78^. Statistical analysis performed using two-tailed paired t-tests. **c,** Representative clonal dynamics of tumor-reactive TCRs visualized by alluvial plots, colored by TfiT-assigned reactivity pre-and post-treatment. **d,** Clonal dynamics across 9 patients with paired pre-and post-treatment samples. Statistically significant changes in clonotype frequency determined by bootstrap-resampling and corrected for multiple comparisons using the Benjamini-Hochberg method. **e,** Box plot of tumor-reactive TCR counts at baseline stratified by clinical response to induction therapy (<VGPR, CR, sCR). All patients with available response and testing data included. **f,** Bar plots of (left) the absolute number of empirically validated tumor-reactive TCRs and (right) the frequency of TfiT-positive BMLs by response group. Statistical comparisons used unpaired t-tests with Welch’s correction. All patients with available response and testing data included.

Together, these data establish a transcriptional blueprint of antitumor T cell responses in hematological malignancies and offer a classifier with translational utility. Because tumor-reactive T cells in hematological malignancies are rare and often phenotypically indistinct from activated bystanders, such a classifier may support enrichment and monitoring of clinically relevant TCRs in both research and therapeutic contexts.

### Pre-existing tumor-reactive T cells correlate with clinical response in NDMM

In solid tumors, efforts to predict anti-tumor immune responses have focused on TCR clonality, neoantigen-specificity, or exhaustion markers - mostly in samples collected post-immunotherapy exposure^10–13,58–61^. By contrast, little is known about how endogenous tumor-specific T cells shape therapeutic outcomes in patients with hematological malignancies.

To address this, we analyzed longitudinal BML samples from patients with newly diagnosed MM (NDMM) undergoing current standard-of-care induction chemotherapy regimens (daratumumab, bortezomib, thalidomide, dexamethasone [Dara-VTd] or bortezomib, lenalidomide and dexamethasone VRD]. We performed paired scRNA/VDJ-seq and bulk TRVα/β-seq on vitally preserved bone marrow samples collected before and after induction therapy (*n* = 15 patients) for direct tracking of tumor-reactive T cell clones using their CDR3 nucleotide sequence as a clonal barcode (**Figure 5a**).

Overall T cell counts remained stable across timepoints (**Figure S12a**), however, post-ASCT samples showed significantly increased clonality (**Figure 5b**), consistent with antigen-driven responses during or after treatment. To assess the behavior of tumor-reactive T cells specifically, we classified all detected TCRs at baseline using the TFiT signature and then tracked their clonal dynamics. Because binomial sampling can bias interpretation of clonal expansion or contraction, we implemented a bootstrapping strategy: 10,000 random draws per patient were used to compute empirical *p*-values for clonal changes across time. Across the cohort, tumor-reactive TCRs were more likely than bystander or orphan clones to persist or significantly expand following therapy (**Figure 5d**). In contrast, virus-specific and orphan TCRs were more frequently lost or remained static.

We next asked whether the abundance of tumor-reactive T cells at diagnosis could predict clinical response to induction therapy. Standard metrics such as the frequency of CD4⁺/CD8⁺/Treg subsets, or prevalence of virus-specific TCRs showed no significant association with therapeutic response (**Figure S12b-c**), in line with prior studies^10^. However, patients who achieved a complete response (CR) or better exhibited a significantly higher frequency of functionally defined tumor-reactive BMLs at baseline, with >21 reactivity-positive cells per functional screen on average^64,65^ (**Figure 5e**). Mirroring this, the abundance of cells classified as tumor-reactive by TFiT in pretreatment bone marrow also correlated with achieving CR (**Figure 5f**).

Together, these data indicate that tumor-reactive TCRs - defined either experimentally or via the TFiT transcriptional signature - are selectively preserved during treatment and correlate with response to induction (immunochemo-)therapy in MM.

### Tumor-reactive T cells underlie responses to immunotherapy in MM and AML

Given the observed association between baseline tumor-reactive T cells and response to induction chemotherapy, we next asked whether this relationship would be further amplified in the context of active immunotherapy - specifically bispecific T cell engagers (bsAbs), which directly depend on the presence of functional, cytotoxic T cells for therapeutic efficacy.

To explore this, we longitudinally profiled bone marrow T cell responses in 18 clinical trial patients with relapsed/refractory multiple myeloma (r/r MM) treated with the BCMA×CD3 bispecific antibody Elranatamab (bsAb)^66,67^. Bone marrow samples were collected at baseline (pre-treatment) and post-cycle 3 (∼90 days), or at relapse, and analyzed by integrated scRNA/VDJ-seq. In total, we profiled 135,567 BMLs across these timepoints and again performed tracking of T cell clonotypes and their antigen-specific states^68^ (**Figure 6a**).

**Figure 6.**
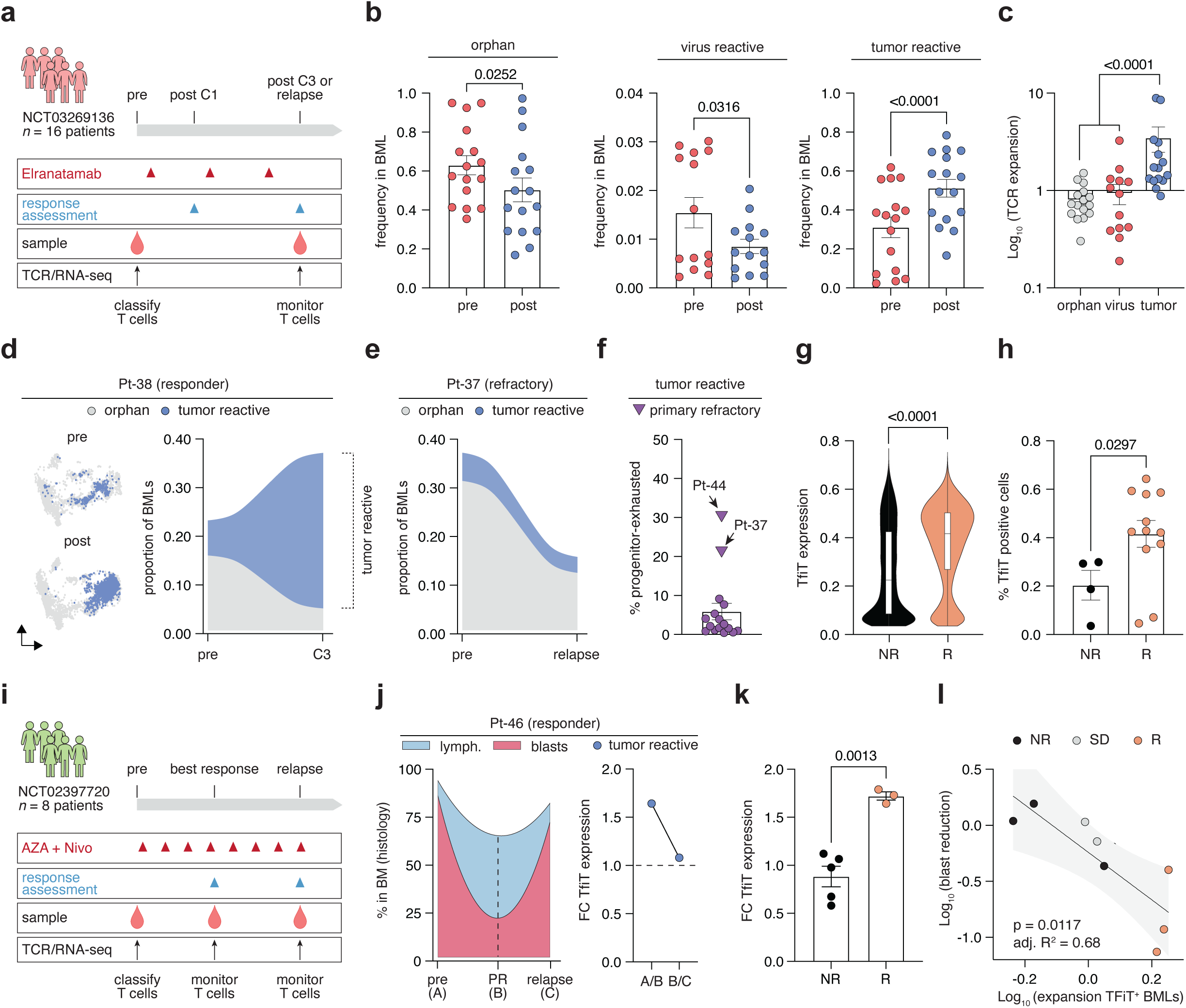
TfiT predicts immunotherapy responses in relapsed/refractory multiple myeloma and AML **a,** Experimental workflow for scRNA-seq and scTCR-seq profiling of 18 relapsed/refractory MM patients pre-and post-treatment treated with the BCMA×CD3 bispecific antibody (bsAb) Elranatamab within the MagnetisMM-1 trial (NCT03269136). **b,** Longitudinal tracking of TCR frequencies stratified by TfiT-predicted reactivity (n=16 patients with available paired samples). Statistical analysis by repeated-measures two-way ANOVA with Tukey’s test. **c,** Expansion of tumor-reactive vs. non-reactive TCRs post-bsAb. Statistical comparison by two-tailed paired t-test. **d-e,** UMAP and alluvial plots from representative clinical responder (Pt-38) and non-responder (Pt-37), visualizing TCR clonotype trajectories pre-and post-bsAb. **f,** Fraction of CD8⁺ tumor-reactive T cells expressing a progenitor-exhausted (PEX) phenotype in primary bsAb-sensitive patients (filled circles) vs. primary bsAb-refractory patients (triangles). **g,** TfiT expression at the single-cell level comparing clinical responders and non-responders after 3 treatment cycles. Statistical comparison by unpaired t-test with Welch’s correction. **h,** Pre-treatment frequency of TfiT⁺ BMLs stratified by clinical response to bsAb after three cycles. Statistical comparison as in (g). **i,** Experimental workflow for scRNA-seq and scTCR-seq profiling of 7 relapsed/refractory AML patients pre-, during-and post-treatment treated with the azacitidine (AZA) + nivolumab (Nivo) within the AARON trial (NCT02397720) **j,** (left): Alluvial plot showing BML and blast dynamics in a representative AML responder (Pt-46) at baseline (A), best response (B) and relapse (C). (right): Fold change in BML TfiT expression between baseline and response (A→B), and response to relapse (B→C) in Pt-46. **k,** Fold change between timepoint A and B in frequency of TfiT⁺ BMLs stratified by clinical response. Statistical comparison by unpaired t-test with Welch’s correction. **l,** Correlation of TfiT⁺ BML expansion and blast reduction across AML patients. Spearman correlation coefficient shown and data points color-coded by patient’s best response.

Classification of TCRs and T cells using the TFiT signature identified tumor-reactive T cells in all r/r MM patients, albeit at variable frequencies. Following bsAb administration, we observed selective expansion of tumor-reactive clones, while orphan and virus-specific TCRs remained largely unchanged (**Figure 6b-c**). We next examined whether these observed clonal dynamics of tumor-reactive TCRs tracked with clinical outcome. In 16 patients with longitudinal sampling, clinical responders showed robust expansion, specifically of tumor-reactive TCRs following bsAb administration (**Figure 6d**). In contrast, non-responders exhibited either stable or contracting tumor-reactive clonotypes (**Figure 6e**), suggesting an impaired capacity for functional T cell recruitment and expansion. These effects were not mirrored by virus-reactive or bystander clones, suggesting the specificity of the response.

Analysis of T cell transcriptional states revealed that CD8⁺ tumor-reactive effector-memory (TR-T_EM_) T cells were the dominant population among expanding clonotypes (**Figure S12d**). By contrast, TR-T_PEX_ cells were rare but appeared to correlate with resistance: both primary refractory patients in the cohort exhibited significantly increased frequencies of baseline TR-T_PEX_ cells (**Figure 6f**). These findings are consistent with emerging evidence that an accumulation of dysfunctional or pre-exhausted T cells in the relapsed myeloma bone marrow may hinder bsAb efficacy^68^. Importantly, a high baseline frequency of TFiT⁺ tumor-reactive T cells correlated with clinical response (**Figure 6h-i**), mirroring our observations in newly diagnosed MM patients treated with induction therapy (**Figure 5f**).

Given the predictive value of tumor-reactive T cells in MM and their association with response to both chemotherapy and bispecific antibodies, we next assessed whether this framework extends to immunotherapy in AML. While allogeneic stem cell transplantation demonstrates that AML is immunologically targetable^69^, clinical responses to immune checkpoint blockade (ICB) have been limited and unpredictable, often attributed to unfavorable T cell phenotypes and low mutational burden^70–72^. To test whether tumor-reactive T cells underlie the rare responses to ICB, we re-analyzed paired scRNA-seq and scTCR-seq data from the phase II AARON trial (NCT02397720)^71^ of azacitidine plus nivolumab in relapsed/refractory (r/r) AML. This cohort comprised 22 longitudinal bone marrow samples from 8 patients, including 3 clinical responders, 3 non-responders, and 2 with stable disease^72^ (**Figure 6i**).

We next applied the TFiT signature to this dataset to determine whether pre-existing tumor-reactive T cells are associated with clinical anti-leukemia response. TFiT-positive clonotypes were detectable at baseline and showed selective expansion in clinical responders following therapy. In contrast, non-responders exhibited minimal or no expansion of tumor-reactive T cells (**Figure 6j**). These dynamics correlated with histological response: patients who achieved partial remission demonstrated a biphasic shift in bone marrow composition: lymphocyte expansion coincided with blast reduction, followed by the reversal of both upon relapse. TFiT expression levels mirrored this trajectory, rising during remission and dropping to baseline at relapse (**Figure 6k**). Across the cohort, the magnitude of tumor-reactive T cell expansion - as inferred by TFiT - correlated with both histological blast reduction and overall clinical response (*R²* = 0.68, *p* = 0.0117), providing further support for the predictive relevance of BMLs with features of tumor reactivity (**Figure 6l**). These findings establish tumor-reactive BMLs as a mechanistic correlate of response to immunotherapy in hematological malignancies and validate the TFiT signature as a broadly applicable tool for their detection. Our approach provides tools for tracking endogenous antitumor immunity across disease states and therapeutic modalities, and offers a proof of concept for using the transcriptional features of rare, but functionally relevant T cell populations to guide patient stratification and immunotherapy development in hematological malignancies.

## Discussion

This study provides a framework for characterizing the nature and specificity of T cell responses in hematological malignancies. We identify a transcriptionally and functionally conserved gene expression program among rare tumor-reactive T cells residing in the bone marrow of patients with two biologically distinct cancers - multiple myeloma (MM) and acute myeloid leukemia (AML). By combining high-sensitivity TCR sequencing, single-cell transcriptomics, and direct identification of HLA-presented antigens, we characterize a discrete subset of bone marrow lymphocytes with features consistent with tumor reactivity, including clonal expansion, polyfunctional cytokine secretion patterns and distinct effector and stem-like gene expression programs. These findings challenge the prevailing notion that hematological malignancies lack robust T cell-mediated immune surveillance and instead suggest that potent tumor-specific T cell responses may be present in principle but remain functionally limited or clinically underappreciated^4,14,73^.

The rarity of tumor-reactive immunological events in hematological malignancies, particularly in treatment-naïve settings, has historically limited their systematic characterization. To address this, we applied a dual strategy combining single-cell functional assays and immunopeptidome-informed expansion assays to identify tumor-reactive clonotypes *in situ* and validate their specificity *in vitro*. This approach enabled us to directly map functionally reactive TCRs to transcriptional states for an integrated view of the phenotype and specificity of tumor-and virus-reactive T cells in MM and AML.

As a result, we found that the tumor-reactive T cell pool in MM and AML shares a conserved transcriptional program, which enables its prospective computational detection through the TFiT signature. This signature captures key phenotypic hallmarks of tumor-reactive T cells, namely, transcriptional convergence, partial effector differentiation, and absence of virus-specific TCR motifs, and outperforms signatures trained on solid tumors in predicting functionally identifiable tumor-reactive clonotypes. Unlike prior work in solid tumors, which often draws from ICB-pretreated samples and distinct resected lesions, our study focuses on untreated patients and the bone marrow microenvironment, which, in part, might explain the observed divergence in T cellular programs^10–13,58–61^.

In relapsed disease, we find that non-response to immunotherapy is characterized by a failure of tumor-reactive T cells to expand or acquire functional phenotypes. This observation supports a model in which exhaustion and TCR-intrinsic features jointly shape therapeutic responsiveness. These findings complement prior evidence from solid tumors suggesting that successful immunotherapy may depend on either recruitment of novel TCRs or reactivation of stem-like T cells within the tumor microenvironment^13^.

Unlike surrogate markers such as tumor mutational burden or clonal hematopoiesis, TFiT provides a prospective, immune-intrinsic measure of anti-tumor immunity anchored in transcriptional phenotype. Its ability to stratify patients treated with checkpoint inhibitors, bispecific antibodies, and immunochemotherapy suggests that transcriptional profiling of bone marrow lymphocytes may serve as both a mechanistic correlate and useful biomarker for clinical outcome.

Our HLA class I immunopeptidomic analysis further reveals a partially shared antigen landscape in MM and AML, including self-derived and noncanonical peptides. Some of these antigens are recognized by tumor-reactive TCRs with convergent motifs, supporting the idea of shared immunogenicity and the existence of public T cell responses targeting hematological malignancies. While neoepitopes derived from somatic mutations were rare, likely reflecting the low mutational burden of these cancers, our data suggest that canonical and noncanonical self-antigens, including cancer-testis antigens and immunoglobulin/BCR-derived peptides, can serve as alternative targets for T cell responses^74,75^ in these malignancies. These findings are consistent with recent work identifying non-mutational immunogenic drivers in myeloid malignancies, such as mis-splicing-derived neoantigens^21^. Our study extends this paradigm by showing that conserved, aberrantly expressed self-antigens can in principle elicit endogenous T cell responses even in the absence of strong mutational signals. However, future studies in larger and more genetically diverse patient populations will be necessary to assess the population-wide prevalence and therapeutic relevance of these epitopes.

From a translational perspective, our approach integrates transcriptional state, antigen presentation, and TCR motif convergence to increase the ‘pre-test’ likelihood of identifying tumor-specific TCRs in cancers with low immunogenicity. This framework could inform the rational design and treatment monitoring of TCR-engineered T cells, bispecific antibodies, or T cell-engaging regimens, particularly in patients with limited mutational targets or poor responses to current therapies.

Several limitations of our study should be noted. First, our functional discovery approach relies on *ex vivo* assays, which may not fully reflect the complexity of the bone marrow microenvironment and are dependent on the functional capacity of the probed T cell. Second, while our workflow is in principle compatible with the detection of HLA class II-restricted epitopes - and we indeed identified several tumor-reactive CD4⁺ T cells - we primarily focused on HLA class I-restricted responses, potentially underrepresenting CD4⁺ T cell-mediated immunity. Third, while shared antigens and TCR motifs suggest potential for public TCR or vaccine approaches, the safety, specificity, and population coverage of these targets require further investigation. Specifically, while our immunopeptidomic analysis identified recurrent nuORF-derived epitopes presented by malignant cells in both MM and AML, the nuORF database used for peptide identification includes entries with low tissue specificity that may, in principle, be expressed in healthy tissues^36^. Recent studies performed in pancreatic ductal adenocarcinoma (PDAC) tissue also detected some nuORF peptides in nonmalignant samples^76^. To address this, we applied stringent peptide spectrum match filtering (FDR <1%) and prioritized peptides recurrently presented across multiple patients but absent in published normal immunopeptidomes. Moreover, transcript-level analysis using GTEx data suggests that several of the genes from which our immunogenic nuORF peptides are derived show negligible expression in hematopoietic or bone marrow tissues^77^. Nonetheless, further safety profiling, including proteomic analysis of matched normal bone marrow and off-target screening, is warranted before therapeutic translation. Finally, the relatively low baseline abundance of tumor-reactive T cells - even in eventual clinical responders - underscores the need for strategies that amplify or sustain these populations *in vivo*.

In summary, we define programs and specificities of tumor-reactive T cells in MM and AML, identify transcriptional correlates of clinical response, and demonstrate the feasibility of detecting rare, but functionally relevant T cells using integrated immunogenomic profiling. These findings support the further development of immunotherapies tailored to the distinct immune biology of hematological malignancies.

## Data and code availability

● Access to all bulk and single-cell WGS/RNA/VDJ sequencing data and annotated computer code required to reproduce the analyses of this manuscript will be available via Zenodo. The repository will be made public upon publication of the peer-reviewed manuscript.
● The original mass spectra, peptide spectrum matches, and the protein sequence database used for searches have been deposited in the public proteomics repository MassIVE (https://massive.ucsd.edu). This dataset will be made public upon publication of the peer-reviewed manuscript.
● All annotated single-cell RNA/V(D)J-sequencing data from newly diagnosed multiple myeloma patients undergoing standard-of-care induction therapy can be accessed at Gene Expression Omnibus (GEO; GSE242883). This dataset will be made public upon acceptance of the manuscript.
● All annotated single-cell RNA/V(D)J-sequencing data from patients enrolled in NCT03269136 can be accessed at Gene Expression Omnibus (GEO; GSE216571). This dataset will be made public upon acceptance of the manuscript.

## Acknowledgements

We thank all members of the Friedrich Lab for their support and fruitful discussions. We are grateful to the DKFZ Sample Processing Lab, the High Throughput Sequencing Unit of the Genomics and Proteomics Core Facility, and the Omics IT and Data Management Core Facility for their technical support. We thank the Clinical Myeloma Registry and Biobank at Heidelberg University Hospital, funded by the Dietmar Hopp Foundation. This study includes samples provided by the NCT Cell and Liquid Biobank (NCT CLB), a member of the Biomaterial Bank Heidelberg (BMBH).

This study was supported by the Deutsche José Carreras Leukämie-Stiftung (DJCLS, PI: M.J. Friedrich; Project-ID DJCLS 01ZI/2022); the Dr. Rolf M. Schwiete Stiftung (PI: M.J. Friedrich; Project-ID 2025-018); the Else Kröner-Fresenius-Stiftung (PI: M.J. Friedrich; Project-ID 2025_EKMS.52); and the Federal Ministry of Education and Research (BMBF) and the Ministry of Science Baden-Württemberg within the framework of the Excellence Strategy of the Federal and State Governments of Germany (PI: M.J. Friedrich; Project-ID ExU 6.1.12);

This work was supported in part by the National Cancer Institute (NCI) Clinical Proteomic Tumor Analysis Consortium (grants U24CA270823 and U01CA271402 to S.A.C.), and the Dr. Miriam and Sheldon G. Adelson Medical Research Foundation (to S.A.C.). K.F. is supported by the Cancer Research UK Early Detection and Diagnosis Programme (C9203). K.Y. is supported by the UCL/UCLH Biomedical Research Centre. B.C. is supported by Cancer Research UK, the Rosetrees Trust, and the UCLH Biomedical Research Centre. M.P. is supported by the European Research Council (ERC Advanced Grant CENTRIC Brain, Project-ID 101141901). M.K. is supported by the German National Academy of Sciences Leopoldina (Project-ID LPDS 2022-13).

N.K. is supported by the International Myeloma Society and the Paula and Rodger Riney Foundation Career Development Award. E.W.G. is supported by the German Ministry of Education and Research (National Center for Tumor Diseases Heidelberg, NCT 3.0 Program ‘Precision Immunotherapy of Brain Tumors’) and the DKTK program (to M.P.). T.B. is supported by the Helmholtz Institute for Translational Oncology Mainz (HI-TRON Mainz) and by a German Cancer Consortium DKTK–DKFZ Joint Funding Project Award (TRUST, to M.P.).

## Author information

### Contributions

T.R.W., N.K. and T.B. designed, and performed experiments, analyzed, and interpreted data. S.St. and N.K. analyzed and interpreted scRNA/VDJ-seq data. M.K., N.K., S.Sc. and M.J.F. performed single-cell experiments. K.F., D.G.-F., E.V., B.C. and K.Y. analyzed longitudinal VDJ-seq data. T.B., K.L., B.S., F.W., C.M.T., and J.L. performed TCR cloning and testing. S.R., L.S.S., J.H.F. and S.Sa. analyzed clinical data. G.M.H., C.C., J.G.A. and S.A.C. performed HLA-I ligandome analyses. S.U. and S.F. performed HLA binding predictions. N.W., E.G., L.B., C.M-T., H.G., K.R., M.S.R., M.P., W.O. and S.B.E. were involved in study design and data interpretation. M.J.F. conceptualized and supervised the study, designed, and performed experiments, analyzed, and interpreted data and wrote the paper with the input from all co-authors.

## Methods

### Human subjects

#### Patients with newly diagnosed multiple myeloma

Bone marrow and peripheral blood samples were obtained from 20 patients at diagnosis (pre-treatment/baseline) and, when available, after induction (immuno-)chemotherapy. All participants provided written informed consent prior to sample collection. The study was conducted in accordance with the principles of the Declaration of Helsinki and the Belmont Report. Ethical approval for the collection, functional testing, and sequencing of human samples was granted by the Ethics Committee of Heidelberg University Medical Faculty (reference numbers S-096/2017, S-777/2024).

#### Patients with newly diagnosed acute myeloid leukemia (AML)

Peripheral blood samples were collected from 7 patients at diagnosis (pre-treatment/baseline). Written informed consent was obtained from all participants, and the study complied with the Declaration of Helsinki and the Belmont Report. Ethical approval for the collection, functional testing, and sequencing of human samples was granted by the Ethics Committee of Heidelberg University Medical Faculty (reference S-744/2024).

#### Patients enrolled in clinical trial NCT02315716^78^

Bone marrow aspirates were collected from 14 patients at baseline (pre-treatment) and at day +100 following autologous stem cell transplantation (ASCT). Written informed consent was obtained from all participants prior to enrollment. The study adhered to the Declaration of Helsinki and the Belmont Report. Ethical approval for the collection, functional testing, and sequencing of human samples was granted by the appropriate UK research ethics committee (reference number 07/Q0502/17).

#### Patients enrolled in clinical trial NCT03269136^66,67^

Samples were collected from 18 patients at diagnosis (pre-treatment/baseline) and either after three cycles of bispecific antibody therapy (post-bsAb cycle 3) or at relapse, whichever occurred first. All patients provided written informed consent for sample collection, tissue sequencing, and review of medical records. The original study was conducted in accordance with the Declaration of Helsinki and the Belmont Report, and was approved by the University of Calgary Institutional Review Board (Ethics ID: HREBA.CC-21-0248) and the resulting dataset re-analyzed for this study.

#### Processing of human bone marrow samples

Bone marrow aspirates were 1:1 diluted in preparation buffer (PBS with 0.1% BSA and 2 mM EDTA), and mononuclear cell separation was performed by density centrifugation (Bicoll separating solution, Biochrom) with diluted bone marrow cells (centrifugation 20 min, 1300g). Cells were carefully aspirated and washed with preparation buffer (centrifugation 5 min at 470g). Red blood cells were lysed using RCL buffer (155 mM NH4Cl, 10 mM KHCO3, 0.1 mM EDTA) for 10 min at room temperature and bone marrow cells were washed (centrifugation 5 min, 470g) and resuspended in preparation buffer. Malignant plasma cells were freshly isolated using CD138 MicroBeads, human (Miltenyi Biotec) according to the manufacturer’s instructions and frozen in 90% FCS (Sigma-Aldrich) supplemented with 10 % DMSO and stored in liquid nitrogen until further use. Non-plasma bone marrow mononuclear cells were frozen after cell counting at 1 × 10^7^ cells per aliquot in 90% FCS (Sigma-Aldrich) supplemented with 10 % DMSO and stored in liquid nitrogen until further use.

#### Cell lines

CD8+ Jurkat reporter cell line was cultured in GlutaMAX-containing RPMI supplemented with 10% heat-inactivated FBS and 1% penicillin-streptomycin. B lymphoblastoid cell lines (B-LCL) were grown in GlutaMAX-containing RMPI 1640 Medium supplemented with 10% FBS, 1% penicillin-streptomycin, 50 mM β-mercaptoethanol, 1 mM sodium pyruvate and 1X MEM Non-Essential Amino Acids, further referred to as B cell media. HEK293T cells were cultured in DMEM supplemented with 10% FBS and 1% penicillin-streptomycin.

#### Antigen-agnostic screening of tumor-reactive T cells

BMLs were isolated as described above. Dead cells were removed using a dead cell removal kit (Miltenyi Biotec). Untouched T cells were isolated using a T Cell isolation kit, human (Miltenyi Biotec) according to the manufacturer’s instructions. Patient-autologous CD138+ plasma or leukemia cells were isolated from thawed bone marrow mononuclear cells using CD138 or CD33 MicroBeads, human (Miltenyi Biotec), respectively. Tumor-reactive T cells were identified in single cell interaction assays on the Lightning™ optofluidic system (Bruker Cellular Analysis). A full clean workflow was carried out on the Lightning system using BLI cleaning solution (PhenomeX) and double distilled water (ddH2O). An OptoSelect™ chip (Bruker Cellular Analysis) was wetted with wetting solution (Bruker Cellular Analysis) according to the manufacturer’s instructions. Human IFNγ, IL-2, and TNFα capture beads (BioLegend) were loaded into the NanoPen™ chambers of the OptoSelect™ chip using opto-electropositioning technology. Subsequently, single T cells and patient matched tumor cells were transferred into each NanoPen™ of the OptoSelect™ chip. As a negative control, one of 12 field of views (FOVs) of the OptoSelect™ chip was loaded with T cells and cytokine capture beads only. As a positive control, a different FOV was loaded with T cells and human T-Activator beads (ThermoFisher Scientific). Cells were co-cultured on the OptoSelect™ chip with cell culture media at 36°C and 5% CO_2_ for 12 h. The OptoSelect™ chip was perfused with a detection antibody mix (72 µl LEGENDplex human Th panel detection antibodies, 4 µl Brilliant Violet 421-anti-hCD137 antibody and 4 µl FITC anti-human CD8 antibody, all from BioLegend) for 30 min. Cells and cytokine capture beads were washed with cell culture medium for 45 min. Subsequently, the OptoSelect™ chip was perfused with a streptavidin-PE solution (8 µl LEGENDplex Streptavidin-PE (BioLegend) in 72 µl PBS) for 30 min. Cells and cytokine capture beads were washed with cell culture medium for 45 min. Fluorescence and brightfield images (DAPI, FITC, CY5, PE, OEP filter) of the cells and cytokine capture beads were acquired at 10x magnification. Fluorescence images were analyzed with Image Analyzer 2.4.16.6 (Bruker Cellular Analysis).

#### Export and TCR sequencing of tumor-reactive T cells

Reactive T cells were unloaded from the respective NanoPens™ of the OptoSelect™ chip using OEP™ and exported into 10 µl of TCL (2X) buffer (Qiagen) in a 96-well PCR plate (Eppendorf). 30 µl of mineral oil (Sigma-Aldrich) was layered on top of each T cell export and cells were centrifuged at 1000*g* for 5 min at RT. RNA was isolated using 20 µl of RNAClean XP magnetic beads (Beckman Coulter). Magnetic beads were washed twice with 80% ethanol (Sigma-Aldrich), dried at RT for 2 min and resuspended in 4 µl of ice-cold RT mix 1 (1.0 µl/well TCRseq primer 1, 1.0 µl/well dNTP mix, 0.2 µl/well RNase inhibitor, 1.8 µl/well ddH_2_O, all from PhenomeX). The export plate was incubated at 72°C for 3 min, followed by a gradual cool to 4°C. 3 µl of ice-cold RT mix 2 (0.5 µl/well TCRseq primer 2, 1.3 µl/well RT buffer, 1.0 µl/well RT additive, 0.1 µl/well RNase inhibitor, 0.1 µl/well RT enzyme, all from PhenomeX) was added to each well of the export plate. First-strand cDNA was synthesized using a thermal cycler (50 min at 42°C, 10 cycles of 2 min at 50°C and 2 min at 42°C, followed by 5 min at 85°C) and purified using 20 µl of MagBind Total Pure NGS magnetic beads (Omega Bio-tek). Magnetic beads were washed twice with 50 µl of 80% ethanol, dried at RT for 2 min and cDNA was eluted by adding 30 µl of nuclease-free water. Agarose gel electrophoresis and DNA quantification (Qubit 4 fluorometer, ThermoFisher Scientific) were performed to verify TCR recovery of each exported T cell. A new PCR plate was prepared on ice (5.0 µl/well PCR enzyme K, 0.4 µl/well TCR amp primer mix (PhenomeX), 3.6 µl/well ddH_2_O, 1.0 µl/well purified cDNA) and V(D)J regions of the TCR alpha and beta chains were amplified using a thermal cycler (3 min at 96°C, 23 cycles of 20 s at 98°C, 30 s at 70°C, 20 s at 72°C, followed by 5 min at 72°C). A reaction mixture, containing unique index pairs for each TCR amplicons, was set up in a new 96-well plate on ice (5.0 µl/well PCR Enzyme K, 0.5 µl/well TCR Index (N7XX and S5XX, Illumina), 4.0 µl/well ddH_2_O, and 0.5 µl/well amplified TCR). Different TCR amplicons were barcoded in an indexing PCR (3 min at 96°C, 10 cycles of 20 s at 98°C, 30 s at 70°C, 20 s at 72°C, followed by 5 min at 72°C). 2 µl of each PCR reaction mixture was pooled and the indexed library was purified using 0.8 volumes of MagBind Total Pure NGS magnetic beads. DNA was washed twice with 80% ethanol; magnetic beads were dried at RT for 3 min and the indexed library was eluted from the beads. The indexed TCR amplicon multiplex was sequenced at a concentration of 10 nM on an Illumina MiSeq system (paired-end 300 bp). TCR V(D)J and CDR3 sequences were identified using MiXCR. TCR sequences with more than 100 reads and productive CDR3 region (no stop codon and in-frame junction) were analyzed. Uncovered sequence regions of full-length TCRα and TCRβ sequences were reconstructed with published TCR sequences exhibiting the highest sequence homology aligned with IMGT/V-QUEST.

### HLA class I immunopeptidomics on CD138+ multiple myeloma cells^37,79–82^

#### Low-input immunoprecipitation of HLA-I:peptide complexes

Immunoprecipitation of HLA-I molecules on tumor cells occurred in three batches. Pt-01 through Pt-08 constituted the first batch, while Pt-09 through Pt-12 constituted the second batch, and Pt-16 to Pt-21 constituted the third batch. Tumor cells isolated from bone marrow aspirates as described above were thawed on ice, lysed in 0.2 mL of lysis buffer at 4°C (20 mM Tris-HCl pH 7.5 (Invitrogen, Waltham, Massachusetts, USA, 15567027), 1 mM Ethylenediaminetetraacetic acid (EDTA) (Invitrogen, Waltham, Massachusetts, USA, 15575-038), 100 mM NaCl (Sigma-Aldrich, St. Louis, Missouri, USA, 71386-1L), 6 mM MgCl_2_ (Sigma-Aldrich, St. Louis, Missouri, USA, 63069-100ML), 60 mM Octyl β-d-glucopyranoside (Sigma-Aldrich, St. Louis, Missouri, USA, O8001-25G), 1 mM phenylmethylsulfonyl fluoride (PMSF) (Sigma-Aldrich, St. Louis, Missouri, USA, 93482-250ML-F), 0.2 mM Iodoacetamide (Thermo Fisher Scientific, Waltham, Massachusetts, USA, A39271), 1.50% Triton X-100 (Sigma-Aldrich, St. Louis, Missouri, USA, T9284-500ML), 1x C0mplete protease inhibitor tablet-EDTA free (Sigma-Aldrich, St. Louis, Missouri, USA, 11873580001), 10 mM NaF (Sigma-Aldrich, St. Louis, Missouri, USA, S7920), 1:100 dilution Phosphatase Inhibitor Cocktail II (Sigma-Aldrich, St. Louis, Missouri, USA, P0044), 1:100 dilution Phosphatase Inhibitor Cocktail III (Sigma-Aldrich, St. Louis, Missouri, USA, P5726), 110 mM Sodium Butyrate (Sigma-Aldrich, St. Louis, Missouri, USA, B5887), 2 µM suberoylanilide hydroxamic acid (SAHA) (Sigma-Aldrich, St. Louis, Missouri, USA, SML0061), 10 mM nicotinamide (Sigma-Aldrich, St. Louis, Missouri, USA, N3376), and 50 μM PR-619 (Lifesensors, Malvern, PA, USA, SI9619: PR-619) in pre-conditioned 0.6 mL Eppendorf tubes, and incubated for 30 min with 0.5 µL benzonase (Sigma-Aldrich, St. Louis, Missouri, USA, E1014-25KU). Eppendorf tubes were pre-conditioned by rinsing twice with 500 µL high-performance liquid chromatography (HPLC) water, incubating overnight with 500 µL HPLC water, and then rinsing twice again with 500 µL HPLC water. After incubating in lysis buffer, the lysates were centrifuged at 15,000 rcf for 20 min at 4°C.

To pre-clear the samples, the supernatants were transferred to another set of pre-conditioned 0.6 mL Eppendorf tubes containing ∼10 µL phosphate buffered saline (PBS)-washed Gammabind Plus Sepharose beads (Sigma-Aldrich, St. Louis, Missouri, USA, GE17-0886-01) and incubated by end-over-end rotation at 4°C for 1 hr. Gammabind Plus Sepharose beads were pelleted at 1500 rcf for 1 min at 4°C. The pre-cleared supernatants were transferred to pre-conditioned 0.6 mL Eppendorf tubes containing PBS-washed Gammabind Plus Sepharose beads (Sigma-Aldrich, St. Louis, Missouri, USA, GE17-0886-01) and 5 µg anti-HLA antibody (W6/32) (Bio X Cell, Lebanon, New Hampshire, USA, BP0079). HLA-I peptide complexes were captured on beads by end-over-end incubation at 4°C for 3 hr. HLA-IP beads were pelleted at 1500 rcf for 1 min at 4°C.

HLA-IP beads were washed on top of a 10 µm PE fritted filter plate (Agilent, Santa Clara, California, USA, S7898A) that was activated with 1 mL acetonitrile (ACN) and equilibrated with 3x 1 mL PBS on a positive pressure manifold (Waters Corporation, Milford, Massachusetts, USA, 186006961). HLA-IP beads were transferred to the activated filter plate with 4°C 1 mL PBS. 500 µL PBS was added to the HLA-IP tube to pick up any remaining beads and transfer them to the filter plate. Beads were washed 2x with 4°C 1 mL wash buffer (20 mM Tris pH 7.5 (Invitrogen, Waltham, Massachusetts, USA, 15567027), 1 mM EDTA (Invitrogen, Waltham, Massachusetts, USA, 15575-038), 100 mM NaCl (Sigma-Aldrich, St. Louis, Missouri, USA, 71386-1L), 60 mM Octyl β-d-glucopyranoside (Sigma-Aldrich, St. Louis, Missouri, USA, O8001-25G), and 0.2 mM Iodoacetamide (Thermo Fisher Scientific, Waltham, Massachusetts, USA A39271)) and 2x with 4°C 1 mL 10 mM Tris-HCl pH 7.5.

#### Low-input HLA-I peptide elution and desalting

All HLA IP samples were acid eluted and desalted in two desalting steps (primary and secondary). For the acid elution and primary desalt of the first batch of samples, Sep-Pak tC18 96-well plate with 40 mg sorbent per well (Waters, Milford, Massachusetts, USA, 186002320) was equilibrated with 2x 1000 µL methanol, 500 µL 99% ACN/0.1% FA, and 4x 1000 µL 1% FA on a positive pressure manifold (Waters Corporation, Milford, Massachusetts, USA, 186006961). Beads were resuspended in 400 µL 3% ACN/5% FA in the filter plate and the filter plate was stacked on top of the equilibrated Sep-Pak tC18 plate. At this step, 50 fmol of Retention Time Standards (iRTs) (JPT Peptide Technologies, Berlin, Germany, RTK-1-10pmol) were spiked into each sample. The HLA-IP beads were washed 2x 200 µL 1% FA. HLA-I peptides were acid eluted from the beads by three, five-minute incubations with 500 µL of 10% acetic acid. The peptides were desalted by 4x 1000 µL 1% FA (after the first FA wash, the filter plate was removed) and eluted from the plate with 250 µL 15% ACN/1% FA and 2x 250 µL 50% ACN/1% FA, frozen at-80°C, and lyophilized.

For the primary desalt of the second batch of samples, a µElution Sep-Pak tC18 96-well plate with 5 mg sorbent per well (Waters Corporation, Milford, Massachusetts, USA, 186002318) was equilibrated with 2x 900 µL methanol, 500 µL 99% ACN/0.1% FA, and 4x 900 µL 1% FA. HLA-IP beads were washed in 200 µL 3% ACN/5% FA in the filter plate which was placed on top of the equilibrated Sep-Pak tC18 plate. At this step, 50 fmol of Retention Time Standards (iRTs) (JPT Peptide Technologies, Berlin, Germany, RTK-1-10pmol) were spiked into each sample. The HLA-IP beads were washed 2x 200 µL 1% FA. HLA-I peptides were acid eluted from the HLA-IP beads by three, five-minute incubations with 500 µL of 10% acetic acid. The peptides were desalted by 4x 900 µL 1% FA (after the first FA wash, the filter plate was removed) and eluted from the plate with 250 µL 15% ACN/1% FA and 2x 250 µL 50% ACN/1% FA, frozen at-80°C, and lyophilized.

The dried HLA-I peptides were stored at-80°C until reduction and alkylation were performed on the second batch of samples. HLA-I peptides were reconstituted in 100 µL 10 mM Tris-HCl pH 7.5 (Invitrogen, Waltham, Massachusetts, USA, 15567027). 2 µL of 250 mM dithiothreitol (DTT) (Thermo Fisher Scientific, Waltham, Massachusetts, USA, A39255) was added to the sample for a final concentration of 5 mM and incubated for 20 min at 50°C while shaking at 1000 rpm. 6 µL of 250 mM iodoacetamide (IAA) (Thermo Fisher Scientific, Waltham, Massachusetts, USA, A39271) was added to the sample for a final concentration of 15 mM and incubated for 30 min at room temperature while shaking at 1000 rpm in the dark. The reaction was quenched with 2 µL of 250 mM DTT, for a final concentration of 5 mM, and incubated for 15 min at room temperature while shaking at 1000 rpm. The sample was brought to 3% ACN/5% FA by adding 90 µL of 6.67% ACN/11% FA.

For all three sets of samples, a secondary desalt was performed on the sample by mirco-scaled basic reversed phase separation on SDB-XC stage tips. Stage tips were set up with two punches of SDB-XC material (CDS analytical, previously Empore 3M, Oxford, Pennsylvania, USA, 98-0604-0223-1). Stage tips were equilibrated with 2x 100 µL methanol, 100 µL 50% ACN/1% FA, and 3x 100 µL 1% FA. The lyophilized peptides were resuspended in 200 µL 3% ACN/5% FA, loaded onto the equilibrated stage tip, and ran through twice to maximize peptide retrieval. HLA-peptides were desalted with 3x 100 µL 1% FA. HLA-peptides were eluted from the SDB-XC stage tips with 60 µL 5% ACN/1% FA, 60 µL 10% ACN/1% FA, and 60 µL 50% ACN/1% FA. Combined elutions were frozen at-80°C, lyophilized, and stored at-80°C^79,80^.

#### LC-MS/MS Analysis

HLA-I immunopeptidome data collection by LC-MS/MS was performed as described previously^85^. The first batch of HLA-I peptides were resuspended in 5 µL 3% ACN/5% FA and 4 µL were injected onto a timsTOF SCP (Bruker, Billerica, Massachusetts, USA). HLA-I peptides were loaded onto a 30 cm analytical column with 1.9 µm ReproSil-Pur C18 silica beads (Dr Maisch HPLC GmbH, Ammerbuch, Germany), packed in-house into a PicoFrit 75 µm diameter fused silica column with a 10 µm emitter (New Objective, Littleton, Massachusetts, PF360-75-10-N-5). HLA-I peptides were eluted using a stepped gradient on a nanoElute LC (Bruker, Billerica, Massachusetts, USA) ranging from 2-15% Solvent B (0.1% FA in ACN) over 60 min, 15-23% over 30 min, 23-35% over 10 min, 35-80% over 10 min and held at 80% for 10 min at 400 nL/min. MS1 scans were acquired from 100-1700 m/z and 1/K0 = 1.70 Vs/cm^2^ to 0.60 Vs/cm^2^ in DDA-PASEF mode. Ten PASEF ramps were acquired with an accumulation and ramp time of 200.0 ms. Precursors above the minimum intensity threshold of 1000 were isolated with 2 Th at <700 m/z or 3 Th at >800 m/z and re-sequenced until a target intensity of 20,000 cts/s followed by a dynamic exclusion of 20 s. The collision energy was lowered linearly as a function of increasing ion mobility from 55.00 eV at 1/K0 = 1.60 Vs/cm^2^ to 15.00 eV at 1/K0 = 0.60 Vs/cm^2^. Standard tryptic precursor isolation polygon placement was optimized for HLA-I peptide species by extending the polygon to include singly charged precursors with >600 m/z^81^.

The second and third batch of samples were acquired on a modified timsTOF SCP, with a higher capacity tims cartridge (Bruker, Billerica, Massachusetts, USA). Peptides were loaded onto a 25 cm Aurora Ultimate CSI nanoflow analytical column with 1.7 µm particle size (IonOpticks, Fitzroy, Victoria, Australia). Peptides were eluted using a stepped gradient on a Vanquish Neo UHPLC System (Thermo Fisher Scientific, Waltham, Massachusetts, USA, VN-S10-A-01). The gradient ranged from 0-15% Solvent B (0.1% FA in ACN) over 60 min, 15-23% over 30 min, 23-35% over 10 min, 35-85% over 10 min and held at 85% for 10 min at 200 nL/min. DDA-PASEF data acquisition parameters are identical to the timsTOF SCP HLA-I parameters described above.

#### HLA-I peptide database search

Raw mass spectra were interpreted with the Spectrum Mill (SM) software package, version 8.01 (Broad Institute; proteomics.broadinstitute.org). Only MS/MS spectra with a precursor sequence MH+ in the range of 700-2000, a precursor charge of +1 to +3, and a minimum of < 5 detected peaks were extracted. Similar spectra with the same precursor m/z acquired in the same chromatographic peak were merged. MS/MS spectra with a sequence tag length >1 (i.e. minimum of three masses separated by the in-chain masses of two amino acids) were searched with no-enzyme specificity. MS/MS spectra were searched against a database comprised of human reference proteome Gencode 42 (ftp.ebi.ac.uk/pub/databases/gencode/Gencode_human/release_34/42) with 50,872 non-redundant protein coding transcript biotypes mapped to the human reference genome GRCh38, 602 common laboratory contaminants, 2043 curated smORFs (lncRNA and uORFs), 237,427 nuORF DB v1.037, and the JPT iRT peptides (JPT Peptide Technologies, Berlin, Germany, RTK-1-10pmol) for a total of 290984 entries^46^. Patient-specific predicted neoepitope peptides (see Methods: NeoEpitope Prediction) were appended to the sequence databases used for searches that are included in the MassIVE deposit. Parameters for SM MS/MS HLA-I search module for the first batch of samples included: ESI-QEXACTIVE-HCD-HLA-v3 30 scoring; fixed modifications: carbamidomethylation of cysteine; variable modifications: protein N-terminal acetylation and deamidation, oxidized methionine, pyroglutamic acid at peptide N-terminal glutamine, deamidation of asparagine, cysteinylation; precursor mass shift range of-18 to 81 Da; precursor mass tolerance of ±15 ppm; product mass tolerance of ±15 ppm; and a minimum matched peak intensity of 40%. For the second batch of samples, parameters for SM MS/MS HLA-I search module differed in: variable modifications: protein N-terminal acetylation and deamidation, oxidized methionine, pyroglutamic acid at peptide N-terminal glutamine, deamidation of asparagine; precursor mass shift range of-18 to 33 Da. Peptide spectrum matches (PSMs) within <1% false discovery rate (% FDR) were confidently assigned for individual spectra via the target decoy estimation of the SM Autovalidation module. PSMs were filtered for precursor charges of +1 to +3, sequence lengths ranging between 8 to 11 amino acids, and a minimum backbone cleavage score (BCS) of 5. BCS is a metric of peptide sequence coverage to enforce uniformly higher minimum sequence coverage for each PSM. The score is a sum after assigning a 1 or 0 between each pair of adjacent amino acids in the sequence (maximum score is peptide length-1) given all ion types, with the goal of decreasing false positive spectra having fragmentation in a limited portion of the peptide by multiple ion types. PSMs were consolidated to peptides through the SM Protein/Peptide Summary module in the file case sensitive mode. A distinct peptide is determined by the highest scoring PSM of a peptide detected for each sample. If different modification states were observed for a peptide, each were reported with a lowercase letter indicating a variable modification (i.e. “C”-carbamidomethylated, “c”-cysteinylated).

#### Subset-specific FDR filtering for nuORFs

Subset-specific FDR was performed as previously described^79^. The aggregate FDR was set to <1% as described above. FDR for the subset of nuORF peptides requires more stringent score thresholding to reach a suitable subset specific FDR of <1%. Subsets of nuORF types were thresholded independently from the HLA dataset through a two-step approach. First, PSM scoring metrics thresholds were tightened on the nuORF subset: minimum SM score of 7, minimum percent scored peak intensity (SPI) of 50%, precursor mass error of ±5 ppm, minimum backbone cleavage score (BCS) of 5. This allows nuORF distributions for each metric to meet or exceed the aggregate distributions. Second, remaining nuORF type subsets with FDR estimate above 1% were further subjected to a grid search to determine the lowest values of BCS and SM score that improved the FDR to <1% for each ORF type in the dataset.

#### Identification of cancer associated antigens (CAAs)

Each HLA-peptide maps to a parent protein, with an associated gene symbol. From preexisting lists of known CAAs^29,80^, the gene symbols of the identified peptides were cross-referenced to identify CAAs in the immunopeptidome.

### Antigen-dependent screening of tumor and virus-specific T cells

A modified version of the MANAFEST assay was used to evaluate T cell responsiveness to tumor and viral antigens^29^. 350,000 tumor cell-depleted BMLs containing T cells from patients Pt-04 to Pt-12 were stimulated *in vitro* with HLA class I-restricted CMV, EBV, Influenza A and Sars-CoV-2 peptide pools (jpt Peptide Technologies) as well as with personalized tumor antigen pools (HLA immunopeptidome-derived; synthesized at >90% purity by GenScript), a shared tumor antigen pool (HLA immunopeptidome-derived; synthesized at >90% purity by GenScript) and without peptide for 10 days. BMLs were cultivated in culture medium (X-VIVO 20 (Lonza) containing 5% of human serum) with 1 µg/ml of each tumor antigen, viral antigen or without peptide (as a reference for non-specific or background clonotypic expansion). On day 3, half of the medium was replaced with fresh medium containing cytokines at a final concentration of 50 IU/ml recombinant human IL-2 (Novartis), 25 ng/ml recombinant human IL-7 and 25 ng/ml recombinant human IL-15 (both from Biolegend). On day 7, half of the medium was replaced with fresh medium containing cytokines at a final concentration of 100 IU/ml IL-2, 25 ng/ml IL-7 and 25 ng/ml IL-15. On day 10, cells were harvested, washed twice with PBS and the cell pellets were snap-frozen. gDNA isolation and UltraDeep TCRB sequencing using the Immunosequencing hsTCRB_v4b workflow were performed by Adaptive Biotechnologies.

### Validation of tumor-reactivity by autologous co-cultures

#### IVT and mRNA production of V(D)J coding gene fragments

V(D)J-coding gene fragments were synthesized by Twist Biosciences. 25 ng gene fragment DNA coding for the V(D)J regions of TCR alpha and beta chains flanked by BsaI restriction sites was mixed with 50 ng of plasmid T7-alpha-TRAC, 50 ng of plasmid T7-beta-TRBC and 10 µl BsaI (New England Biolabs) and Golden Gate Reaction Mix was incubated in a thermocycler (30 cycles of 1 min at 42°C, 1 min at 16°C, followed by 5 min at 60°C). 0.5 µl of the reaction mixture was incubated with 1 µl of 1:10 diluted forward primer, 1 µl of 1:10 diluted reverse primer, 5 µl of Clone Amp™ HiFi PCR Premix (ThermoFisher Scientific) and 3 µl of ddH2O in a thermocycler (30 s at 98°C, 30 cycles of 5 s 98°C, 5 s 62°C, 10 s at 72°C, followed by 30 s at 72°C). The ligated DNA fragment was purified using 1.8 volumes of MagBind Total Pure NGS magnetic beads (Omega Bio-tek) and eluted in 10 µl of ddH2O. The ligation of the DNA fragment was verified by Sanger sequencing (Eurofins Genomics) and 1 µg of DNA was *in vitro* transcribed using the T7 mScript™ standard mRNA production system (Cellscript). 5’-capped, 3’-polyadenylated mRNA was purified using 1.8 volumes of RNAClean XP magnetic beads (Beckman Coulter) and eluted in 40 µl ddH2O.

#### Rapid T cell expansion for tumor specificity validation

PBMC were thawed and 0.5×10^6^ CD8+ T cells were sorted using a CD8+ T cell isolation kit, human (Miltenyi Biotec). CD8+ T cells were co-cultured with feeder cells (10×10^7^ PBMCs irradiated at 40 Gy with Gammacell 1000 Elite irradiator) in X-VIVO™ 15 hematopoietic cell media supplemented with 2% human AB serum, 300 U/ml recombinant human IL-2 (Novartis Pharmaceuticals) and an initial concentration of 30 ng/ml CD3 antibody (OKT3, ThermoFisher Scientific) in T25 cell culture flasks (Cellstars). Half of the cell culture media was replenished with fresh media 3 times a week. At day 14 post start of co-culture, T cells were used for electroporation with IVT-transcribed mRNA of respective TCRA and TCRB chains to be tested.

#### mRNA transfection and functional testing of transgenic TCR-expressing T cells

2×10^6^ expanded T cells in 20 µL nucleofector P3 solution (Lonza) were transfected with 1 µg of TCR coding mRNA in each well of a 96-well unit using an 4D-Nucleofector™ electroporator (EO-115 program, Lonza). Cells were rested for 10 min at RT, 180 µl of per-warmed TexMACS™ Medium (Miltenyi Biotec) was added and electroporated cells were incubated for 24 h in a 48-well plate. 50,000 electroporated autologous T cells and 50,000 tumor cells were co-cultured in 200 µl cell culture media in a 96-well U-bottom plate for 16 h at 37°C. As controls, TCR-transgenic T cells were cultured with autologous CD8-depleted PBMCs, 1 µg/ml CEFT peptide pool (JPT Peptide Technologies), or alone. As positive control, T cells were stimulated with human T cell TransAct™ (Miltenyi Biotech) according to the manufacturer’s instructions. In addition, all former conditions were tested for mock electroporated autologous T cells (without TCR encoding mRNA). Flu-specific TCR mRNA-transfected autologous CD8+ T cells were co-cultured with tumor cells and the aforementioned controls. Cells were washed twice with staining buffer (DPBS, 3% FBS, 2 mM EDTA), incubated with human Fc block™ (BD Biosciences) for 10 min at RT and stained with CD8-FITC, mTRBC-PE, CD69-PE-Cy7 and CD137-APC antibodies (all from BioLegend) for 20 min on ice in the dark. A 1:1000 diluted live/dead blue viability dye (Life Technologies) was used. Compensation was performed using OneComp eBeads™ (ThermoFisher Scientific). Fixed cells were acquired on the BD Canto II (BD Biosciences) running BD FACSDiva Software. Data were analyzed using FlowJo (v.10.8.1; Tree Star).

### Epitope mapping of tumor-reactive TCRs

#### B cell isolation and EBV immortalization to generate B-LCLs

PBMCs were thawed and diluted to 1 x 10^7^ cells/ml in B cell isolation buffer (PBS+2 % FCS+ 1 mM EDTA). B cell isolation was performed by negative selection with the EasySep Human B cell Isolation Kit (17954, StemCell Technologies) following manufacturer’s guidelines. Isolated B cells were counted and diluted in B cell media to 0.3 x 10^6^ cells/ml. Epstein-Barr virus-containing supernatant from B95-8 cells is added to the cell suspension to reach a final cell density of 0.25 x 10^6^ B cells/ml. EBV-induced immortalization was further supported by holo-transferrin (30 µg/ml, T0665, Sigma-Aldrich) and CpG ODN 2006 (2.5 µg/ml, tlrl-2006, InvivoGen). B cells were plated in round-bottom 96-well plates at a cell density of 5 x 10^4^ per well and expanded for subsequent analyses.

#### Generation of TCR-transgenic Jurkat cell lines

Fully humanized TCRs were ordered as custom synthesis from Twist Bioscience in a third-generation pLVX-EF1α-IRES-Puro lentiviral expression vector (631253, Takara). HEK293T were cotransfected with a VSV-G envelope expression plasmid (pMD2.G) and a packaging plasmid (psPAX2) following the TransIT-Virusgen Transfection (MIR6700, Mirus) guidelines. Viral supernatants were harvested 48 hours post transfection and filtered supernatants were added onto Jurkat reporter T cells. The following day viral supernatant was replenished with fresh T cell media and selection of TCR-transgenic Jurkat T cells with puromycin (1 µg/ml) was started 96 hours after transduction.

#### Epitope mapping assays and analysis

Co-cultures were performed with the T-FINDER platform as previously described (citation T-Finder). In brief, TCR-transgenic Jurkat T cells were plated with patient-derived B-LCLs in a 3:1 ratio in 96-well U-bottom plates. Peptides were synthesized at GenScript (>90% purity) and added in a final concentration of 10 µM. Co-cultures and T cell activation were analyzed after 16 hours. Staining solution was prepared in flow cytometry buffer (PBS, 2 % FCS and 2 mM EDTA) and the following antibodies were diluted 1:200: anti-human CD2 PerCP-Cy5.5 (RPA-2.10, 300216, Lot: B399713, B382499), anti-human CD20 PE (2H7, 302306, Lot: B380281), anti-human CD69 APC-Cy7 (FN50, 310912, Lot: B260870), anti-human CD3 PE-Cy7 (UCHT1, 300420, Lot: B368934) and anti-HA.11 APC (16B12, 901524, Lot: B362222). All samples were acquired on a SonyID7000 spectral analyzer and data was analyzed using FlowJo (10.6.1). The proportion of CD69-positive cells was determined by calculating the integrated area under the histogram that did not overlap with control samples, employing FlowJo’s population comparison (Overton) technique.

#### Antigen HLA promiscuity testing using tetramers

The following HLA tetramers were used: Flex-T™ HLA-A*01:01 Monomer UVX, 280001, Biolegend; Flex-T™ HLA-A*02:01 Monomer UVX, 280004, Biolegend; Flex-T™ HLA-A*03:01 Monomer UVX, 280005, Biolegend; Flex-T™ HLA-A*11:01 Monomer UVX, 280007, Biolegend. Tetramer:peptide complexes were prepared according to the manufacturer’s instructions. 20 µl Flex-T™ monomer UVX (200 µg/ml, Biolegend) was mixed with 20 µl peptide solution (500 µM, synthesized at >95% purity by GenScript Biotech) in PBS. Monomers were illuminated with UV light (365 nm, 8 Watts, 5 cm) for 30 min on ice. Monomers were subsequently incubated for 30 min at 37°C in the dark. 4.4 µl of PE-streptavidin or APC-streptavidin solution (0.2 mg/ml, Biolegend) were added, mixed and incubated for 30 min on ice in the dark. A blocking solution was prepared by mixing 1.6 µl 50 mM D-Biotin (Biolegend) and 192.4 µl PBS. After incubation of HLA-matched BML samples with tetramers, 2.4 µl of blocking solution was added and tetramers were incubated on ice for at least 30 min in the dark. Cells were washed twice with staining buffer (PBS, 3% FBS, 2 mM EDTA), incubated with human Fc block™ (BD Biosciences) for 10 min at RT and stained with 100 µl 1:100 diluted Flex-T™ complex for 30 min on ice in the dark. Cells were co-stained with anti-CD3-Pacific Blue™ and anti-CD8-FITC antibodies (both from Biolegend) as well as 1:1000 diluted live/dead™ fixable yellow dye (Life Technologies) for 30 min on ice in the dark. Compensation was performed using OneComp eBeads™ (ThermoFisher Scientific). Fixed cells were acquired on the BD Canto II (BD Biosciences) running BD FACSDiva Software. Data were analyzed using FlowJo (v.10.8.1; Tree Star).

### Single-cell transcriptome / V(D)J and CITE-seq analyses

#### Single-cell sequencing and data preprocessing

For single-cell RNA (scRNA), V(D)J, and CITE-seq analyses, bone marrow lymphocytes (BMLs) were counted and divided into eight aliquots per patient sample. Each aliquot was stained with TotalSeq™ hashtag antibodies (BioLegend) and fluorescently labeled antibodies prior to pooling. Viable CD45⁺CD3⁺ cells were isolated using fluorescence-activated cell sorting on a BD Aria Fusion™ system (**Figure S1a**). Single-cell capture, reverse transcription, and library preparation were performed on the Chromium platform (10x Genomics) using the Single Cell 5′ Reagent Kit v2, following the manufacturer’s protocol, with 40,000 cells loaded per channel. Library quality was assessed, and final libraries were paired-end sequenced (26 and 92 bp) on an Illumina NovaSeq 6000 S2 lane. Raw sequencing data encompassing gene expression, V(D)J, and cell hashing information were processed using Cell Ranger (v7.1) with the ‘multi’ command, referencing the GRCh38 genome (2020-A) and V(D)J reference (v7.1.0).

#### Quality control and normalization

Demultiplexing of single-cell data was performed using the HTODemux function in Seurat (v5.0.1)^83^, classifying cells as singlets, doublets, or unassigned based on hashtag oligos.

Heterotypic doublets were identified with the scDblFinder package and, along with hashtag-identified doublets, removed from further analysis. Cells lacking a productive alpha/beta TCR were excluded. All datasets were normalized using SCTransform^84,85^ and mapped to a reference dataset of pre-annotated bone marrow scRNA-seq profiles from newly diagnosed MM patients. This step facilitated removal of contaminating CD3⁻ cells, likely arising from sorting artifacts. Mapping was performed using the MapQuery function in Seurat, following the SCTransform v2 vignette (https://satijalab.org/seurat/articles/sctransform_v2_vignette.html). Only cells with >200 detected genes, <10% mitochondrial content, and identified as singlet CD3⁺ T cells were classified as high-quality and retained for analysis.

#### scVDJ analysis

TCR clonotypes were defined based on the CDR3 amino acid sequences of paired alpha and beta chains as annotated by Cell Ranger. In cases where multiple alpha or beta chains were detected, the top-expressed chain was retained. Unique TCRs were identified per patient and assigned stable IDs across analyses. Diversity indices were computed in R using the vegan package (v2.6-4)( https://vegandevs.github.io/), and clonotype proportions were calculated within each post-QC dataset per patient.

#### Back-projection of antigen-specific TCRs to single-cell transcriptomes

Tumor-reactive TCRs identified via functional screening were mapped to single-cell RNA/TCR datasets based on exact CDR3 amino acid sequence matches. When a match was observed for both TRA and TRB - or for TRB alone - the single-cell clonotype was annotated as tumor-reactive. This approach identified 62 unique tumor-reactive clonotypes, comprising a total of 2,133 single cells across all patients. TCRs expanded in MANA-stimulated conditions were identified using ImmunoSeq UltraDeep sequencing (Adaptive Biotechnologies) and analyzed using the FEST tool^29^ (http://www.stat-apps.onc.jhmi.edu/FEST/) with an odds ratio threshold of 2, assuming 250,000 cells per well. TCRs with >1% abundance at baseline were excluded. Expanded TCRs were back-projected onto single-cell datasets via exact CDR3 TRB amino acid matching. Publicly annotated virus-and tumor-reactive TCRs were mapped by matching TRB CDR3 sequences to the VDJdb^86^ (downloaded January 25, 2024) and McPAS-TCR^87^ (downloaded February 1, 2024) databases, respectively. Tumor-reactive TCRs targeting TP53 and RAS were included based on literature from Levin et al. (2021)^88^, Kim et al. (2022)^89^, and Malekzadeh et al. (2019)^90^. HLA-matched TCRs were filtered and mapped to single-cell populations according to their restriction: HLA class I TCRs were assigned to CD8/NKT cells, and HLA class II TCRs to CD4⁺ T cells. Ambiguously annotated TCRs (e.g., with dual virus/tumor specificity) were flagged in all outputs.

#### Module scores of T cell dysfunction and cytotoxicity^31^

The following module scores were added to single cell dataset using Seurat’s AddModuleScore(…, assay= “RNA”):

*T cell dysfunction score:*

“LAG3”,“HAVCR2”,“PDCD1”,“PTMS”,“FAM3C”,“IFNG”,“AKAP5”,“CD7”,“PHLDA1”,“ENT

PD1“,”SNAP47“,“TNS3”,“CXCL13”,“RDH10”,“DGKH”,“KIR2DL4”,“LYST”,“MIR155”,“RAB

27A“,”CSF1“,”CTLA4“,”TNFRSF9“,”CD27“,”CCL3“,”ITGAE“,”PAG1“,”TNFRSF1B“,”GALNT1“,”GBP2“,”MYO7A“

*T cell cytotoxicity score:*

“FGFBP2”,“CX3CR1”,“FCGR3A”,“S1PR5”,“PLAC8”,“FGR”,“C1orf21”,“SPON2”,“CD300A”,“

TGFBR3“,”PLEK“,”S1PR1“,”EFHD2“,”KLRF1“,”FAM65B“,”C1orf162“,”STK38“,”SORL1“,”F

CRL6“,”TRDC“,”EMP3“,”CCND3“,”KLRB1“,”SAMD3“,”ARL4C“,”IL7R“,”GNLY“

#### Clustering of tumor-reactive cells

To cluster the validated tumor-reactive T cells separately, we first isolated cells that were annotated with tumor-reactive TCRs. These cells were then integrated afresh using Harmony version 0.1.1^91^ and Seurat^83^, with the integration performed on a per-patient basis using default settings. The top 15 PCA dimensions, calculated based on the 2,500 most variable features (with TCR variable genes excluded), were used for this integration. Clusters were determined based on the top 15 Harmony dimensions at a resolution setting of 0.5, merged and annotated manually based on marker gene expression. The expression of features within these clusters was depicted using density plots, facilitated by the Nebulosa package version 1.8.0^92^.

### Differential gene expression and signature derivation

Differential gene expression (DGE) analysis was conducted using the presto package (v1.0.0; https://github.com/immunogenomics/presto) via the wilcoxauc() function on raw assay data. TCR variable genes were excluded from the DGE analysis. Visualizations were generated using the pheatmap package (v1.0.12) with scaled average gene expression values (calculated via Seurat’s AverageExpression() function), or Seurat-native plotting functions.

To derive the TFiT (Tumor-reactive Feature-inferred T cell) signature, we performed DGE in the establishment cohort after downsampling each TCR reactivity group to a maximum of 2,000 cells. Tumor-reactive cells were defined based on experimental evidence, literature-based annotation, and the MANA assay, and were compared to cells annotated as virus-reactive (CMV, EBV, Influenza A, SARS-CoV-2) or orphan T cells. Marker genes were filtered using thresholds of adjusted p-value < 0.01 and log2 fold-change > 0.5.

In the full cohort, a similar DGE workflow was performed with reactivity group sizes capped at 5,000 cells. Eight iterations of gene signature candidates were assessed, prioritizing genes ranked by AUC values. The final TFiT signature comprised the top 15 genes meeting the above statistical criteria and demonstrating robust performance across datasets.

To assess the predictive value of TFiT and other published gene signatures, we conducted gene set enrichment analysis (GSEA) using AUCell (v1.24.0)^93^, following the method described in Lowery et al. (2022)^12^. Area under the ROC curve (AUROC) metrics were computed with the pROC package (v1.18.5)^94^. Tumor-reactive cells were assigned as the positive class (1), and all others as negative (0). AUCell enrichment scores were used as predictors in ROC analyses. The optimal threshold for the TFiT score was determined using the coords() function to achieve sensitivity and specificity > 0.8 in the validation cohort (AUC: 0.8953; threshold: 0.48, specificity: 0.801, sensitivity: 0.807). In the establishment cohort, this threshold yielded an AUC of 0.9795, specificity of 0.938, and sensitivity of 0.952. This threshold (0.48) was applied throughout the study for classifying tumor-reactive T cells based on their gene expression signature.

#### CDR3 similarity and network analysis

After the identification of tumor-reactive and the annotation of virus-reactive TCRs as described above, the sequences from all different classes of TCRs were extracted from TCR and scRNA/VDJ sequencing data were used for similarity analyses. All analyses were implemented in custom R functions. TCR sequences of non-annotated clonotypes from scRNA/VDJ were used as bystander-TCRs. CDR3 TRA and TRB sequence similarities were evaluated using local alignment with BLOSUM45 and a gap opening cost of 10 as described previously^95–97^ utilizing the msa package in R. To normalize local alignment scores, they were divided by the self-alignment score of the query-sequence. The separated scaled similarity was aggregated by calculating a geometric mean for scaled TRA and TRB similarity for each pairwise comparison. A bootstrapping methodology was employed to establish a similarity score significance threshold. In this method all pairwise comparisons of non-annotated TRA-TRB pairs were calculated using the same approach outlined above. This resulted in a background distribution from over 100.000 TCR sequences from TCR-clones found in the bone marrow. Two thresholds at either a 95% or 99% percentile of these background distributions were calculated. The normalized similarities of virus-reactive TCRs were analyzed with the same approach. Across patients, similarities of tumor-reactive TCR TRA-TRB were computed. Additionally, gliph^53^ as implemented in the R package turbogliph (https://github.com/HetzDra/turboGliph) was used to identify clusters of TRA or TRB sequences. Gliph-based clusters were calculated using the turbogliph function independently for TRA and TRB chains of the included clonotypes. Visualization was rendered through heatmaps filtering values above the 95% bootstrapping limit using the ComplexHeatmap R package^98^. To show that gliph^53^ was able to identify similarities in either TRA or TRB, each TRA or TRB identified gliph-cluster is shown together with the pairwise similarity analysis. A graphical representation of similarities filtered with the 99% threshold was created with the igraph^99^ and ggraph (https://ggraph.data-imaginist.com/) R packages. Distinct clusters of TRA-TRB similarity had their sequence motifs for TRA and TRB illustrated using the msa^100^ and ggseqlogo^101^ R packages.

### Analysis of longitudinal newly diagnosed MM samples during induction therapy

Paired scRNA-seq and scVDJ-seq data from n=9 patients before and after induction treatment were used to trace clonotype dynamics. Clonal overlap was defined by matching paired TRA and TRB CDR3 sequences. Relative abundance of each clonotype was calculated per timepoint. To assess statistical significance of clonal dynamics, we applied a bootstrapping-based empirical cumulative distribution function (ECDF) analysis, as previously described^68^. This involved 10,000 resampling iterations of the TCR repertoire at each timepoint. ECDF-derived p-values were adjusted for multiple comparisons using the Benjamini-Hochberg method.

Clonotype dynamics were categorized into five groups: expansion (significant increase), contraction (significant decrease), stable, complete loss (absent post-treatment), and replacement (only detected post-treatment).

### Analysis of longitudinal relapsed/refractory MM samples during bispecific antibody therapy^68^

Bispecific antibody (bsAb)-treated scRNA-seq datasets^68^ were mapped to the same pre-annotated reference of bone marrow T cells from newly diagnosed MM patients. Only cells with productive TCRs were retained. GSEA was performed using AUCell ^93^(v1.24.0) with the previously defined TFiT signature and fixed enrichment thresholds.

Virus-reactive TCRs were annotated via exact TRB CDR3 sequence matching to VDJdb^86^, without HLA filtering. TCR overlap between timepoints was assessed based on full-length CDR3 amino acid identity. Clonotypes with >40% of their cells scoring above the TFiT threshold across timepoints were designated as reactive. Alluvial plots were generated using the ggalluvial^102^ (v0.12.5) package. Patients who relapsed before the end of cycle 3 were designated as clinical non-responders / bsAb-refractory.

### Analysis of longitudinal r/r AML samples during AZA/nivolumab therapy^71,72^

AML patient datasets^72^ (kindly provided by Hussein A. Abbas) were mapped to the same pre-annotated T cell reference from newly diagnosed MM samples. Integration was performed using Seurat^83,91^’s IntegrateLayers(…, method = HarmonyIntegration) with log-normalized counts. Dimensionality reduction was performed using UMAP (RunUMAP(…, reduction = “harmony”, dims = 1:30)).

GSEA of the TFiT signature was conducted using AUCell with previously established thresholds. To correlate immune response with clinical parameters, TFiT dynamics and AML burden were evaluated between timepoints A (pre-treatment) and B (post-initial exposure). Fold-changes in average TFiT expression and blast percentages were log₁₀-transformed and compared via linear regression using R’s lm() function.

### Deep TCRB sequencing of clinical trial samples

Genomic DNA was isolated from peripheral blood samples using QIAamp DNA isolation kit (QIAGEN) as per the manufacturer’s instructions. TCR beta chain (TCRB) deep sequencing was performed on purified DNA from isolated bone marrow or blood mononuclear cells to detect rearranged TCRβ gene sequences using hsTCRB Kit (Adaptive Biotechnologies) according to the manufacturer’s protocol. The prepared library was sequenced on an Illumina MiSeq. TCR identification, error correction and CDR3 was performed using the *Decombinator* pipeline^103^, available at https://github.com/innate2adaptive/Decombinator. Samples with < 2000 total TCR counts were removed. TCR α and β-chains were aggregated to clonotypes for each donor by amino acid CDR3 sequence. Clonality was defined as the inverse of diversity calculated using the Shannon diversity index and Rényi entropy as implemented by the *vegan* package (https://cran.r-project.org/web/packages/vegan/index.html). To account for differences in sample size, a weighted subsampled of 2000 counts was taken 100 times for each sample as the mean diversity calculated.

### Whole-genome sequencing of tumor cells and germline controls

WGS was performed on an Illumina NovaSeq 6000 instrument with S4 flow cells in paired-end mode (2×151 bp). Previously described protocols were used for the extraction of nucleic acids, library preparation, and computational processing^104^.

### Prediction of immunoglobulin-derived neoepitopes in multiple myeloma

To identify putative immunoglobulin-derived neoepitopes in MM, we developed a multistep bioinformatic workflow integrating somatic variant calling, HLA typing, and peptide binding prediction. Somatic variants were identified from tumor-enriched CD138⁺ MM cells by comparing with matched germline data from PBMCs for exclusion of germline-encoded or non-somatic alterations. Variants located in non-coding regions were removed from downstream analyses. HLA class I haplotypes were inferred using OptiType^105^ (v1.3.1), and candidate neoepitopes were predicted from non-synonymous SNVs and indels using NetMHCpan^106^ (v4.1). To focus on HLA-presented ligands, only 8-and 9-mer peptides with a predicted eluted ligand rank ≤2% were retained. Variant allele frequency (VAF) was used to prioritize mutations with high clonal representation, minimizing artifacts due to tumor heterogeneity or subclonality. To mitigate the risk of selecting peptides with off-target reactivity, we filtered candidate neoepitopes against the UniProtKB^107^ protein database using PeptideMatch^108^ (v1.0). Peptides with sequence identity to known human proteins were excluded as potential self-antigens. The final list of predicted neoepitopes was ranked based on HLA-binding affinity and VAF of the underlying mutation, yielding a high-confidence set of immunoglobulin-derived neoantigen candidates for functional screening and validation.

### Quantification and statistical analysis

Data are represented as individual values or as mean ± SEM, as indicated. Group sizes (n) and applied statistical tests are indicated in figure legends. Statistical significance and multiple hypothesis testing corrections were assessed as indicated in figure legends. All reported p values are two-tailed. All analyses were performed using either R v4.3.2 (www.R-project.org) or GraphPad Prism 10.0. For functional experiments, bone marrow samples were blinded to the experiment performers. Shannon and all others indices were calculated using diversity() vegan R package or Gini() as stated in methods above.

Due to the nature of this study, sample size determination was not applicable, as all available samples were included in this study. All cells passing QC (**Figure S1-2**, methods) were included in downstream analyses on a single-cell basis.

### Data visualization

Tabular data from single-cell sequencing analyses above were processed using the tidyverse suite of packages [https://CRAN.R-project.org/package=tidyverse] and visualized in the R programming environment using the ggplot2 package or the Python programming environment using the matplotlib package. Data from all other analyses were visualized using GraphPad Prism 10.0. Figures were produced using Adobe Illustrator 2024.

