## Supplementary Materials for "Conserved programs and specificities of T cells targeting hematological malignancies"

**The PDF file includes:**

Supplementary Figures S1-S12

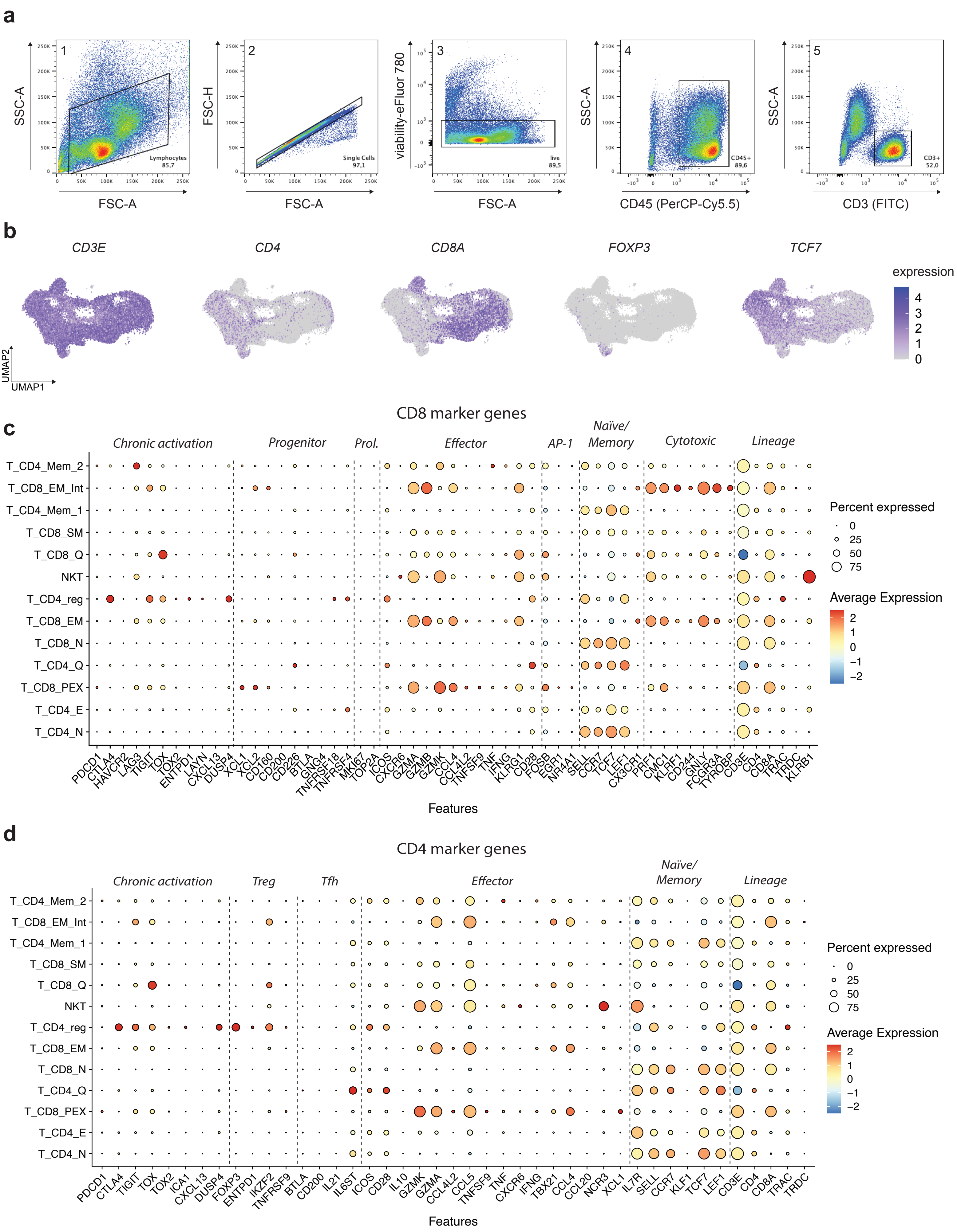

a
SSC-A
FSC-A
Lymphocytes 85,7
FSC-H
Single Cells 97,1
viability-eFluor 780
live 89,5
CD45 (PerCP-Cy5.5)
CD45+ 89,6
CD3 (FITC)
CD3+ 52,0
b
CD3E
CD4
CD8A
FOXP3
TCF7
expression
UMAP2
UMAP1
c
CD8 marker genes
Chronic activation
Progenitor
Prol.
Effector
AP-1
Naïve/ Memory
Cytotoxic
Lineage
T_CD4_Mem_2
T_CD8_EM_Int
T_CD4_Mem_1
T_CD8_SM
T_CD8_Q
NKT
T_CD4_reg
T_CD8_EM
T_CD8_N
T_CD4_Q
T_CD8_PEX
T_CD4_E
T_CD4_N
PDCD1 CTLA4 HAVCR2 LAG3 TIGIT TOX TOX2 ENTPD1 LAYN CXCL13 DUSP4 XCL1 XCL2 CD160 CD200 CD226 BTLA GNG4 TNFRSF18 TNFRSF4 MKI67 TOP2A ICOS CXCR6 GZMA GZMB GZMK CCL4 CCL4L2 TNFSF9 TNF IFNG KLRG1 CD28 FOSB EGR1 NR4A1 SELL CCR7 TCF7 LEF1 CX3CR1 PRF1 CMC1 KLRF1 CD244 GNLY FCGR3A TYROBP CD3E CD4 CD8A TRAC TRDC KLRB1
Features
Percent expressed
Average Expression
d
CD4 marker genes
Chronic activation
Treg
Tfh
Effector
Naïve/ Memory
Lineage
PDCD1 CTLA4 TIGIT TOX TOX2 ICA1 CXCL13 DUSP4 FOXP3 ENTPD1 IKZF2 TNFRSF9 BTLA CD200 IL21 IL6ST ICOS CD28 IL10 GZMK GZMA CCL4L2 CCL5 TNFSF9 TNF CXCR6 IFNG TBX21 CCL4 CCL20 NCR3 XCL1 IL7R SELL CCR7 KLF1 TCF7 LEF1 CD3E CD4 CD8A TRAC TRDC
Features
Percent expressed
Average Expression

**Figure S1. Single-cell profiling of bone marrow lymphocytes (BMLs) in newly diagnosed multiple myeloma.**

**a,** Representative fluorescence-activated cell sorting (FACS) gating strategy for isolation of $CD45^{+}CD3^{+}$ T cells from bone marrow aspirates. Sorted populations were processed for single-cell 5' RNA-seq, CITE-seq, and paired TCR V(D)J profiling using the 10x Genomics Chromium platform.
**b,** UMAP embedding of BMLs from all patients. Overlaid surface protein expression levels from CITE-seq used to annotate canonical T cell populations.
**c, d,** Dot plots displaying average expression and detection frequency of canonical marker genes across $CD8^{+}$ (c) and $CD4^{+}$ (d) T cell clusters. Marker sets were curated based on previously defined signatures[103–105].

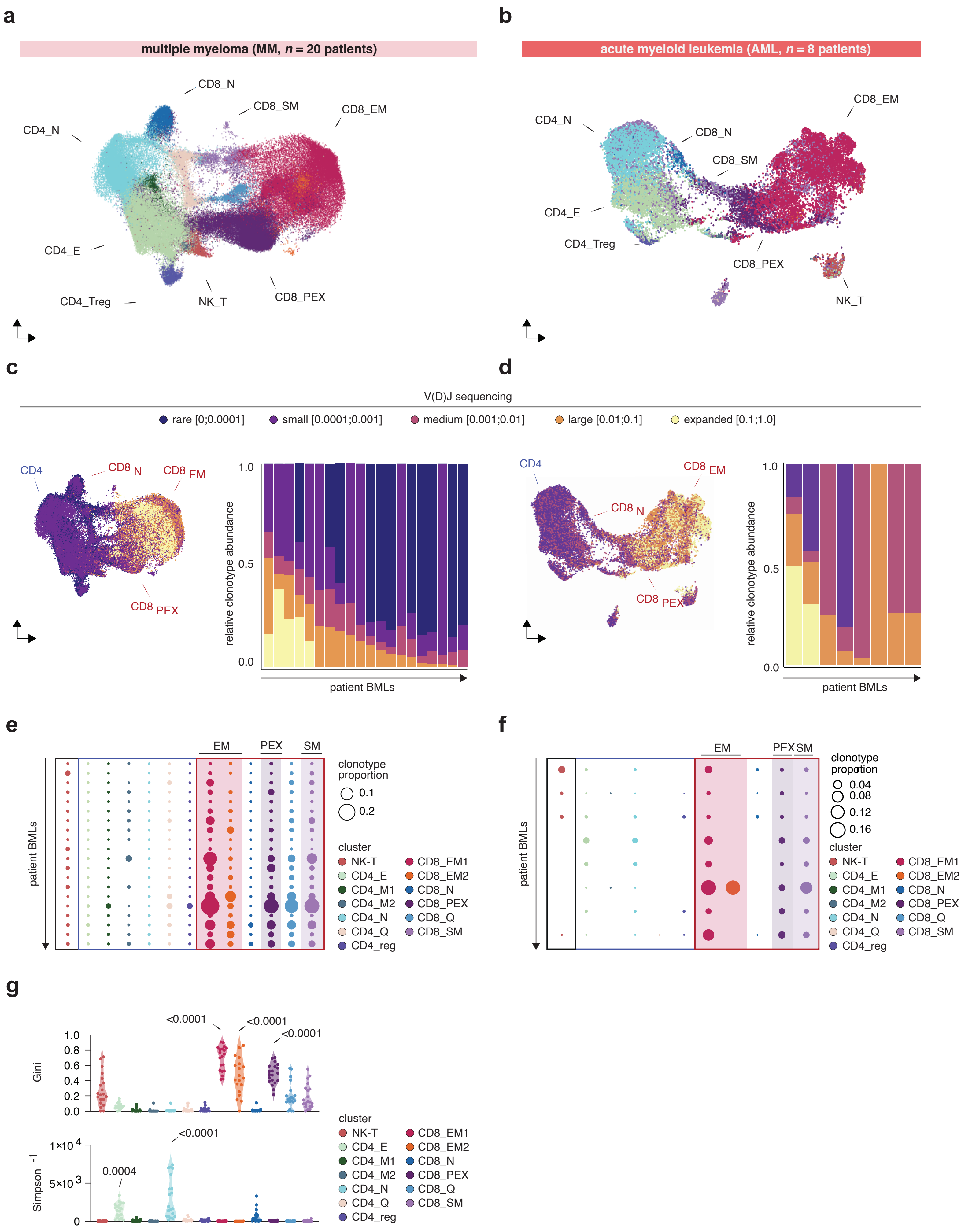
a
multiple myeloma (MM, n = 20 patients)
CD8_N
CD8_SM
CD8_EM
CD4_N
CD4_E
CD4_Treg
NK_T
CD8_PEX
b
acute myeloid leukemia (AML, n = 8 patients)
CD8_EM
CD4_N
CD8_N
CD8_SM
CD4_E
CD4_Treg
CD8_PEX
NK_T
c
d
V(D)J sequencing
rare [0;0.0001]
small [0.0001;0.001]
medium [0.001;0.01]
large [0.01;0.1]
expanded [0.1;1.0]
CD4
CD8 N
CD8 EM
CD8 PEX
relative clonotype abundance
1.0
0.5
0.0
patient BMLs
e
f
EM
PEX
SM
patient BMLs
clonotype proportion
0.1
0.2
0.04
0.08
0.12
0.16
cluster
NK-T
CD4_E
CD4_M1
CD4_M2
CD4_N
CD4_Q
CD4_reg
CD8_EM1
CD8_EM2
CD8_N
CD8_PEX
CD8_Q
CD8_SM
g
Gini
1.0
0.8
0.6
0.4
0.2
0.0
<0.0001
<0.0001
<0.0001
Simpson -1
1×10 4
5×10 3
0
0.0004
<0.0001

**Figure S2. Comparative analysis of BML TCR landscapes in newly diagnosed multiple myeloma (NDMM) and acute myeloid leukemia (AML).**

**a,** UMAP projection of T cell subtypes with productive TCRs in the NDMM cohort (N=20 patients; n=187,015 cells; n=132,501 TCRs passing QC).
**b**, UMAP projection of T cell subtypes with productive TCRs in the NDMM cohort (N=8; n=21,714 cells; n=11,302 TCRs passing QC).
**c,** (Left) Clonotype expansion mapped onto the UMAP and categorized by relative abundance (expanded: 0.1-1.0; large: 0.01-0.1; medium: 0.001-0.01; small: 0.0001-0.001; rare: <0.0001). (Right) Relative proportions of expansion categories across patients in NDMM cohort.
**d,** (Left) Clonotype expansion mapped onto the UMAP and categorized by relative abundance (N=8 patients)(expanded: 0.1-1.0; large: 0.01-0.1; medium: 0.001-0.01; small: 0.0001-0.001; rare: <0.0001). (Right) Relative proportions of expansion categories across patients. For AML cohort[66] in timepoint A.
**e,** Average proportion of cells per TCR clonotype across annotated T cell subsets and patients in NDMM cohort(dot size scaled to clonotype abundance).
**f,** Average proportion of cells per TCR clonotype across annotated T cell subsets and patients in AML cohort[66] (dot size scaled to clonotype abundance).
**g,** Gini index and inverse Simpson index plotted per transcriptionally defined T cell cluster to assess clonal dominance and diversity, respectively (N=20 patients).

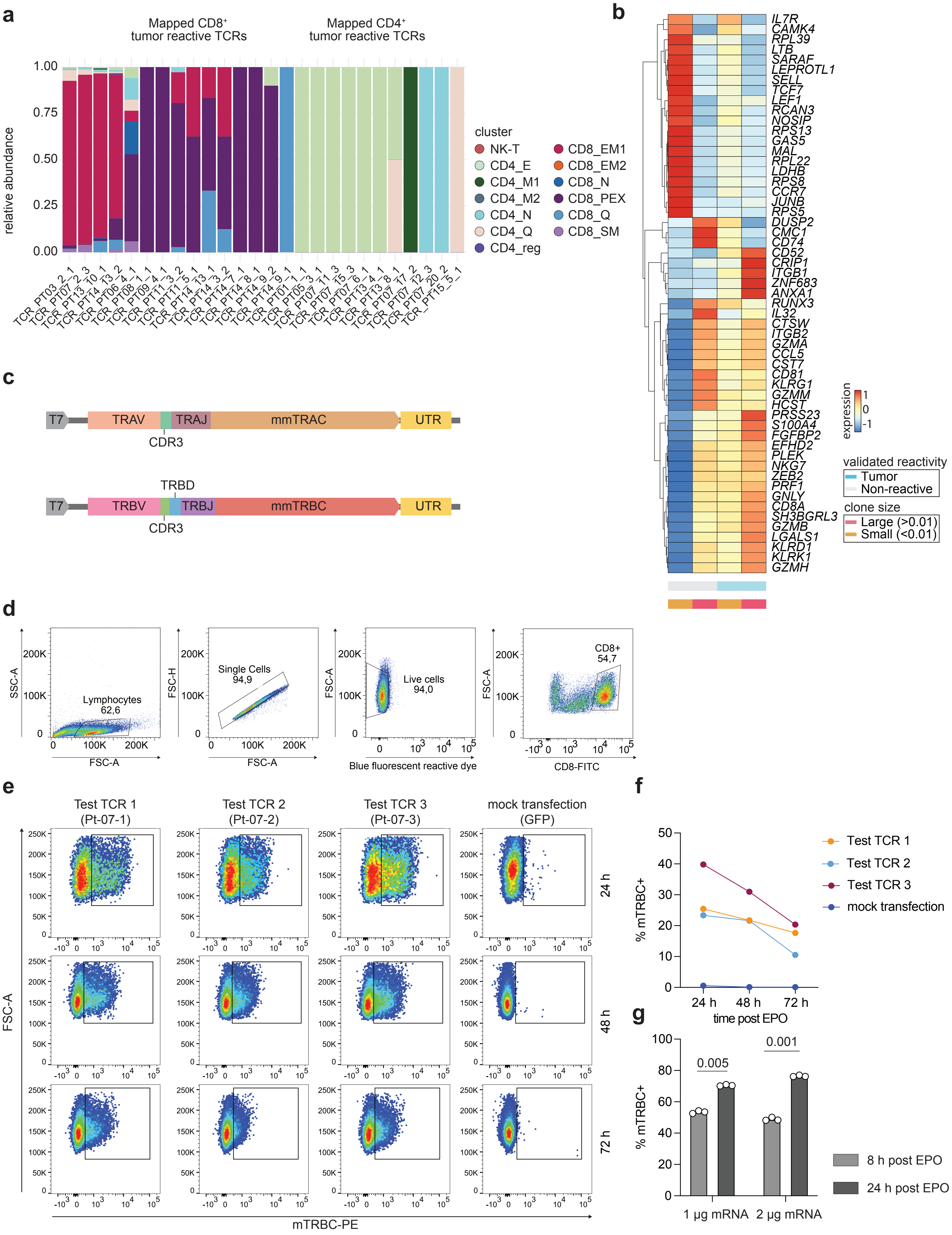

a
Mapped CD8+ tumor reactive TCRs
Mapped CD4+ tumor reactive TCRs
relative abundance
1.00
0.75
0.50
0.25
0.00
cluster
NK-T
CD4_E
CD4_M1
CD4_M2
CD4_N
CD4_Q
CD4_reg
CD8_EM1
CD8_EM2
CD8_N
CD8_PEX
CD8_Q
CD8_SM
TCR_PT03_2_1
TCR_PT07_2_3
TCR_PT13_10_1
TCR_PT14_13_2
TCR_PT06_4_1
TCR_PT08_1_1
TCR_PT09_4_1
TCR_PT11_3_2
TCR_PT11_5_1
TCR_PT14_13_1
TCR_PT14_3_2
TCR_PT14_7_1
TCR_PT14_8_1
TCR_PT14_9_2
TCR_PT14_9_1
TCR_PT01_1_1
TCR_PT05_3_1
TCR_PT07_11_3
TCR_PT07_15_3
TCR_PT07_6_1
TCR_PT13_4_1
TCR_PT13_8_1
TCR_PT07_17_2
TCR_PT07_12_3
TCR_PT07_20_2
TCR_PT15_5_1
b
IL7R
CAMK4
RPL39
LTB
SARAF
LEPROTL1
SELL
TCF7
LEF1
RCAN3
NOSIP
RPS13
GAS5
MAL
RPL22
LDHB
RPS8
CCR7
JUNB
RPS5
DUSP2
CMC1
CD74
CD52
CRIP1
ITGB1
ZNF683
ANXA1
RUNX3
IL32
CTSW
ITGB2
GZMA
CCL5
CST7
CD81
KLRG1
GZMM
HCST
PRSS23
S100A4
FGFBP2
EFHD2
PLEK
NKG7
ZEB2
PRF1
GNLY
CD8A
SH3BGRL3
GZMB
LGALS1
KLRD1
KLRK1
GZMH
expression
1
0
-1
validated reactivity
Tumor
Non-reactive
clone size
Large (>0.01)
Small (<0.01)
c
T7
TRAV
TRAJ
mmTRAC
UTR
CDR3
TRBD
T7
TRBV
TRBJ
mmTRBC
UTR
CDR3
d
SSC-A
Lymphocytes
62,6
FSC-A
FSC-H
Single Cells
94,9
FSC-A
FSC-A
Live cells
94,0
Blue fluorescent reactive dye
FSC-A
CD8+
54,7
CD8-FITC
e
Test TCR 1
(Pt-07-1)
Test TCR 2
(Pt-07-2)
Test TCR 3
(Pt-07-3)
mock transfection
(GFP)
24 h
48 h
72 h
FSC-A
mTRBC-PE
f
% mTRBC+
Test TCR 1
Test TCR 2
Test TCR 3
mock transfection
24 h
48 h
72 h
time post EPO
g
% mTRBC+
0.005
0.001
1 µg mRNA
2 µg mRNA
8 h post EPO
24 h post EPO

**Figure S3. Transcriptional and functional characteristics of tumor-reactive TCR clonotypes.**

**a,** Transcriptional cluster distribution of functionally validated TCR clonotypes, matched to $CD4^+$ or $CD8^+$ T cells by integrated scRNA-seq, V(D)J data.
**b,** Heatmap showing mean expression of top 20 marker genes across large (>1% of cells) and small (<1%) tumor-reactive and non-reactive TCR clonotypes.
**c,** Schematic of linear DNA constructs used for IVT of patient-derived TCRA/B V(D)J regions fused to murine TRAC or TRBC constant regions to enable flow cytometric detection.
**d,** Gating strategy to identify T cells expressing electroporated transgenic TCRs by surface detection of murine TRBC (mTRBC).
**e,** Representative flow cytometry plots showing mTRBC expression in transgenic T cells.
**f, g,** Surface expression kinetics of TCRs following electroporation, including time-course (f) and mRNA dose titration (g). Statistical significance was evaluated using one-way analysis of variance (ANOVA), followed by Tukey's post hoc test for multiple comparisons.

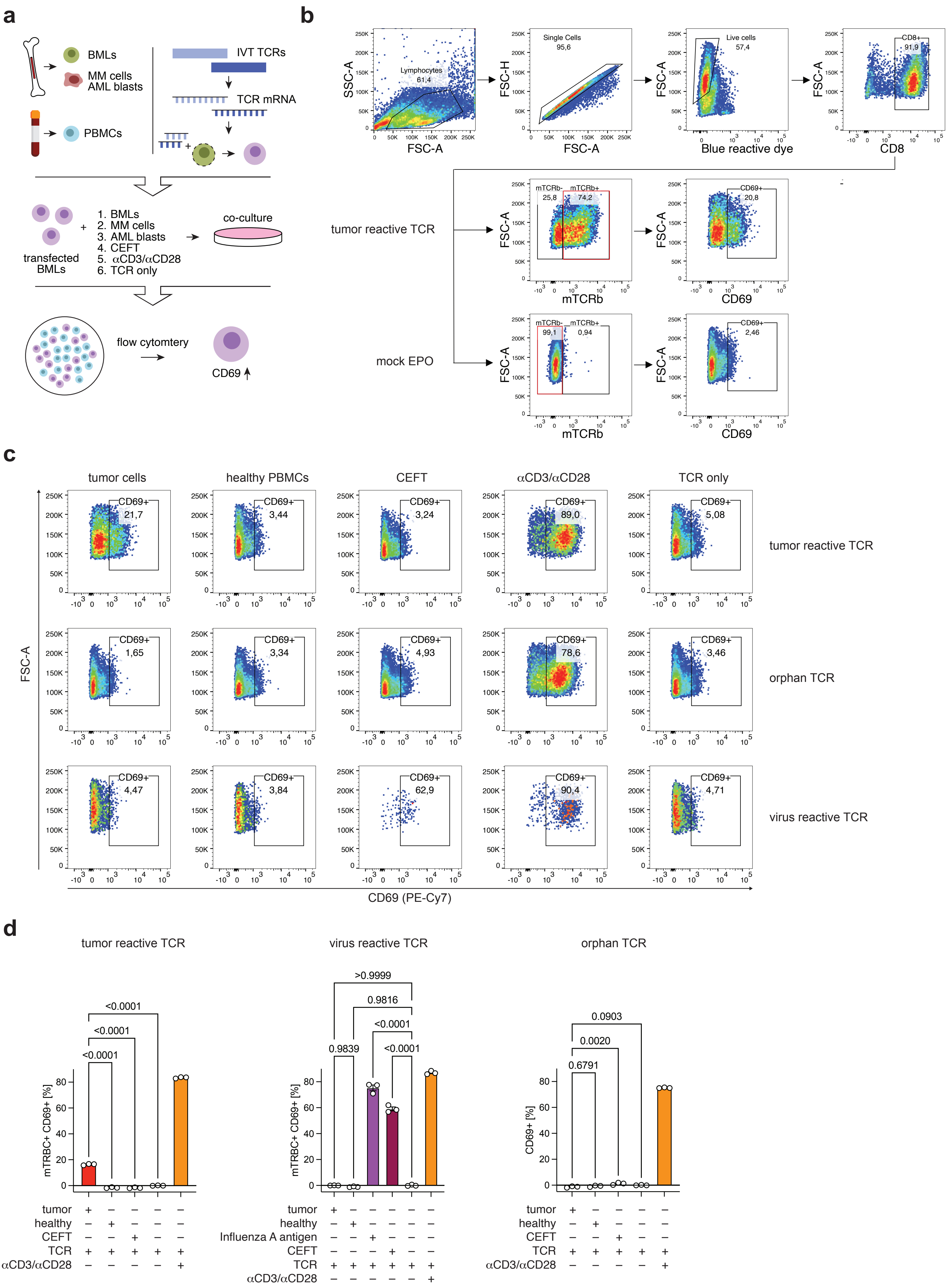

a
BMLs
MM cells
AML blasts
PBMCs
IVT TCRs
TCR mRNA
1. BMLs
2. MM cells
3. AML blasts
4. CEFT
5. αCD3/αCD28
6. TCR only
transfected BMLs
co-culture
flow cytomtery
CD69
b
SSC-A
FSC-A
FSC-H
Lymphocytes 61,4
Single Cells 95,6
Live cells 57,4
Blue reactive dye
CD8+ 91,9
CD8
tumor reactive TCR
mock EPO
mTCRb- 25,8
mTCRb+ 74,2
CD69+ 20,8
mTCRb- 99,1
mTCRb+ 0,94
CD69+ 2,46
mTCRb
CD69
c
tumor cells
healthy PBMCs
CEFT
αCD3/αCD28
TCR only
CD69+ 21,7
CD69+ 3,44
CD69+ 3,24
CD69+ 89,0
CD69+ 5,08
tumor reactive TCR
CD69+ 1,65
CD69+ 3,34
CD69+ 4,93
CD69+ 78,6
CD69+ 3,46
orphan TCR
CD69+ 4,47
CD69+ 3,84
CD69+ 62,9
CD69+ 90,4
CD69+ 4,71
virus reactive TCR
FSC-A
CD69 (PE-Cy7)
d
tumor reactive TCR
virus reactive TCR
orphan TCR
mTRBC+ CD69+ [%]
CD69+ [%]
<0.0001
<0.0001
<0.0001
>0.9999
0.9816
<0.0001
0.9839
<0.0001
0.0903
0.0020
0.6791
tumor
healthy
CEFT
Influenza A antigen
TCR
αCD3/αCD28

**Figure S4. Patient-autologous functional testing of tumor-reactive TCRs.**

**a,** Schematic of the TCR validation workflow using autologous T cells and tumor cells.
**b,** Gating strategy to detect transgenic TCR (mTRBC) expression and CD69 as an early activation marker.
**c,** Representative CD69 expression for a functionally validated tumor-reactive TCR (top), an orphan (non-reactive) TCR (middle), and a virus-reactive TCR (bottom) under co-culture with autologous tumor cells or controls.
**d,** Quantification of CD69 upregulation across three independent experiments (electroporation + co-culture). Statistical analysis was performed using repeated-measures one-way ANOVA followed by Tukey's multiple comparisons test.

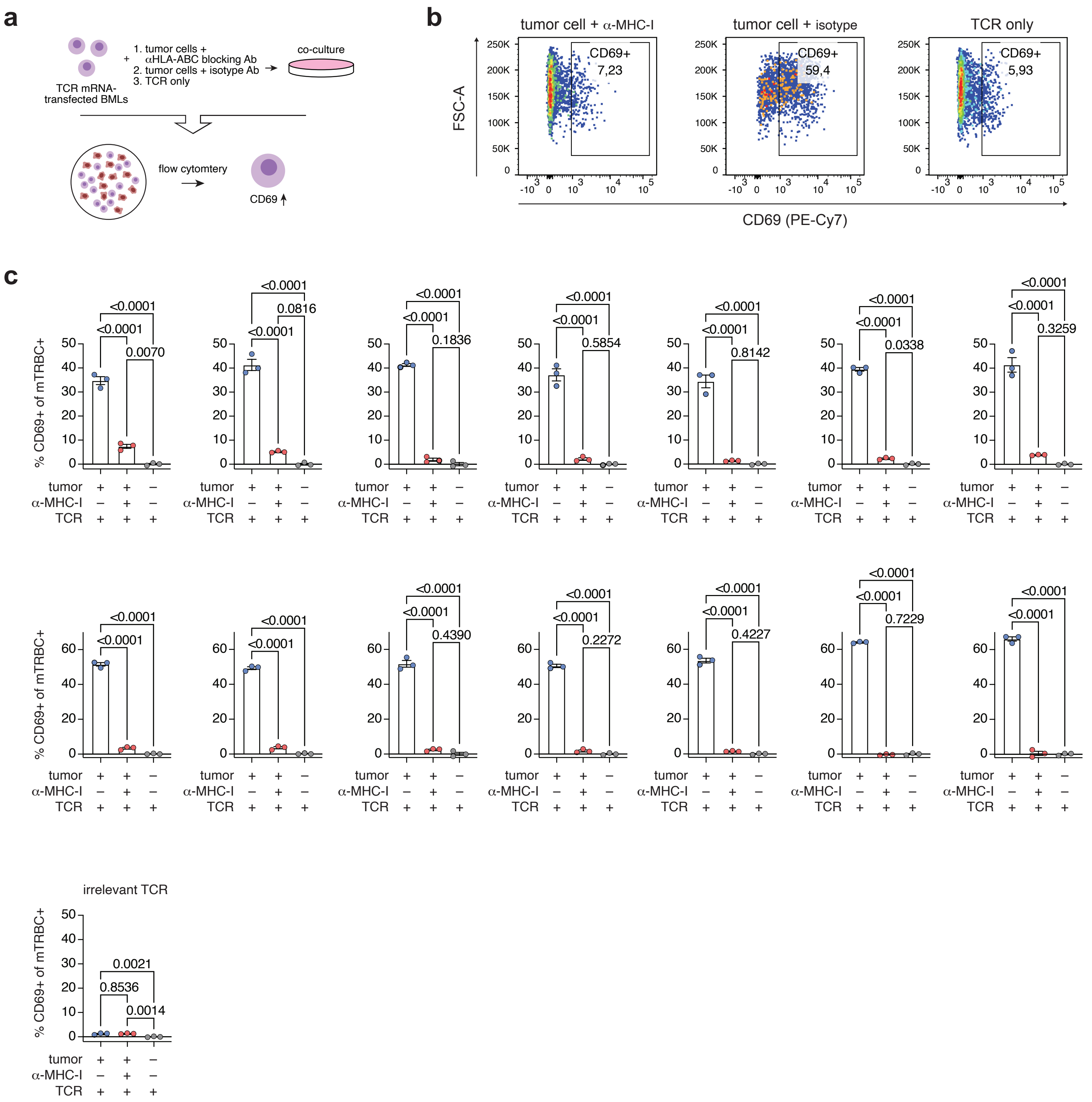
a
1. tumor cells + αHLA-ABC blocking Ab
2. tumor cells + isotype Ab
3. TCR only
co-culture
TCR mRNA-transfected BMLs
flow cytomtery
CD69
b
tumor cell + α-MHC-I
tumor cell + isotype
TCR only
CD69+ 7,23
CD69+ 59,4
CD69+ 5,93
FSC-A
CD69 (PE-Cy7)
c
% CD69+ of mTRBC+
tumor
α-MHC-I
TCR
<0.0001
0.0070
0.0816
0.1836
0.5854
0.8142
0.0338
0.3259
0.4390
0.2272
0.4227
0.7229
irrelevant TCR
0.0021
0.8536
0.0014

**Figure S5. MHC dependence of tumor-reactive TCR activation.**

**a,** Experimental schematic showing co-culture of TCR-transfected T cells with autologous tumor cells in the presence of anti-HLA-ABC blocking antibody or isotype control.
**b,** Representative flow cytometry plots from TCR-transfected T cells under the indicated conditions.
**c,** Quantification of CD69 upregulation in 14 tumor-reactive TCRs. An orphan TCR was used as a negative control. Statistical analysis was conducted using repeated-measures one-way ANOVA with Tukey's test for post hoc multiple comparison adjustment.

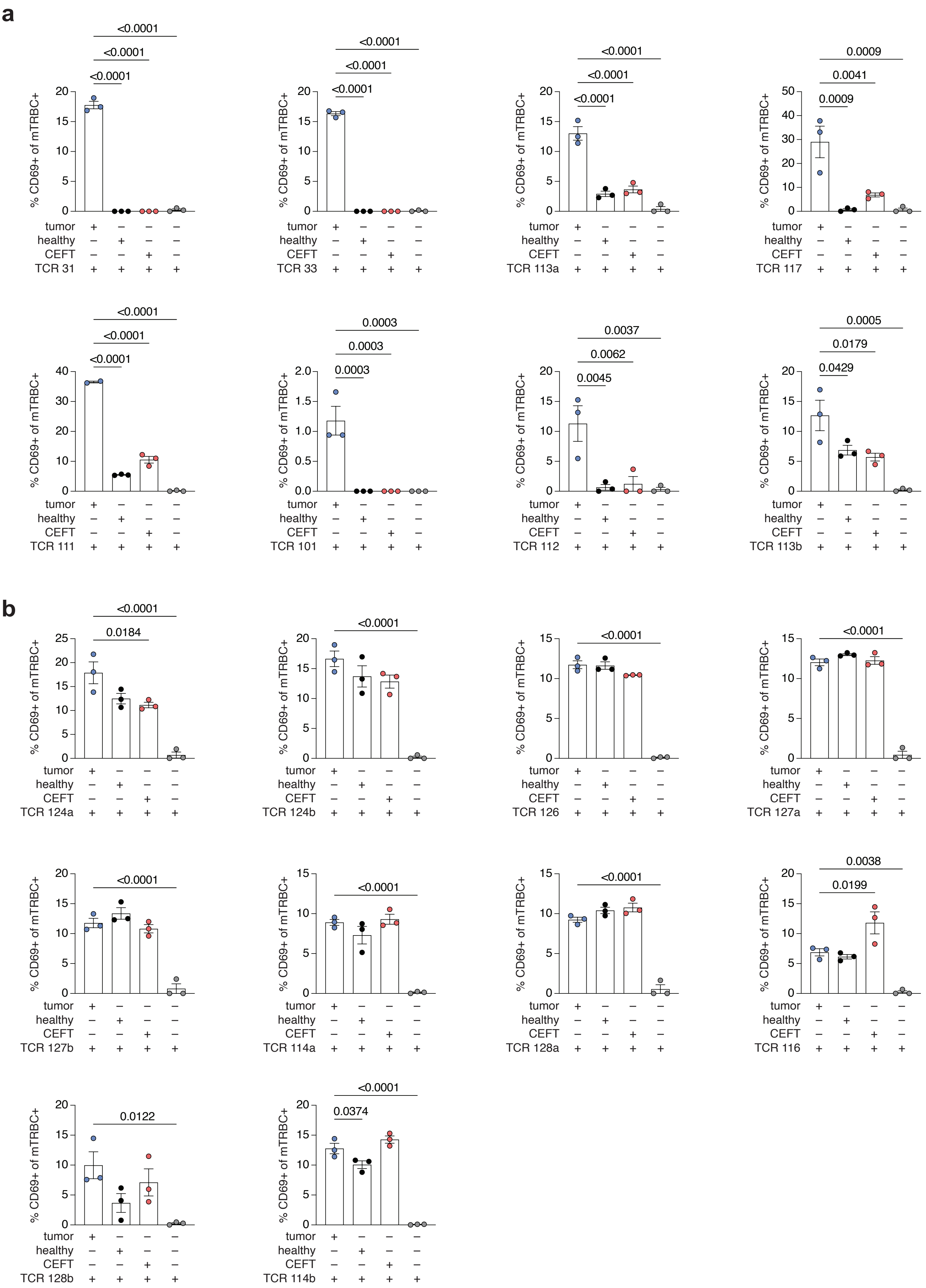
a
<0.0001
<0.0001
<0.0001
% CD69+ of mTRBC+
tumor
healthy
CEFT
TCR 31
TCR 33
TCR 113a
0.0009
0.0041
0.0009
TCR 117
TCR 111
0.0003
0.0003
0.0003
TCR 101
0.0037
0.0062
0.0045
TCR 112
0.0005
0.0179
0.0429
TCR 113b
b
0.0184
TCR 124a
TCR 124b
TCR 126
TCR 127a
TCR 127b
TCR 114a
TCR 128a
0.0038
0.0199
TCR 116
0.0122
TCR 128b
0.0374
TCR 114b

**Figure S6. Specificity profiling of tumor-reactive and cross-reactive TCRs.**

**a, b,** TCR-transfected autologous T cells were co-cultured with autologous tumor cells, PBMCs (healthy control), CEFT peptide pool (viral control), or no antigen. CD69 expression was measured after 16 hours. Data are shown for (a) tumor-reactive and (b) cross-reactive TCRs, with three replicate experiments per condition. Statistical comparisons were performed using repeated-measures one-way ANOVA followed by Tukey's post hoc test.

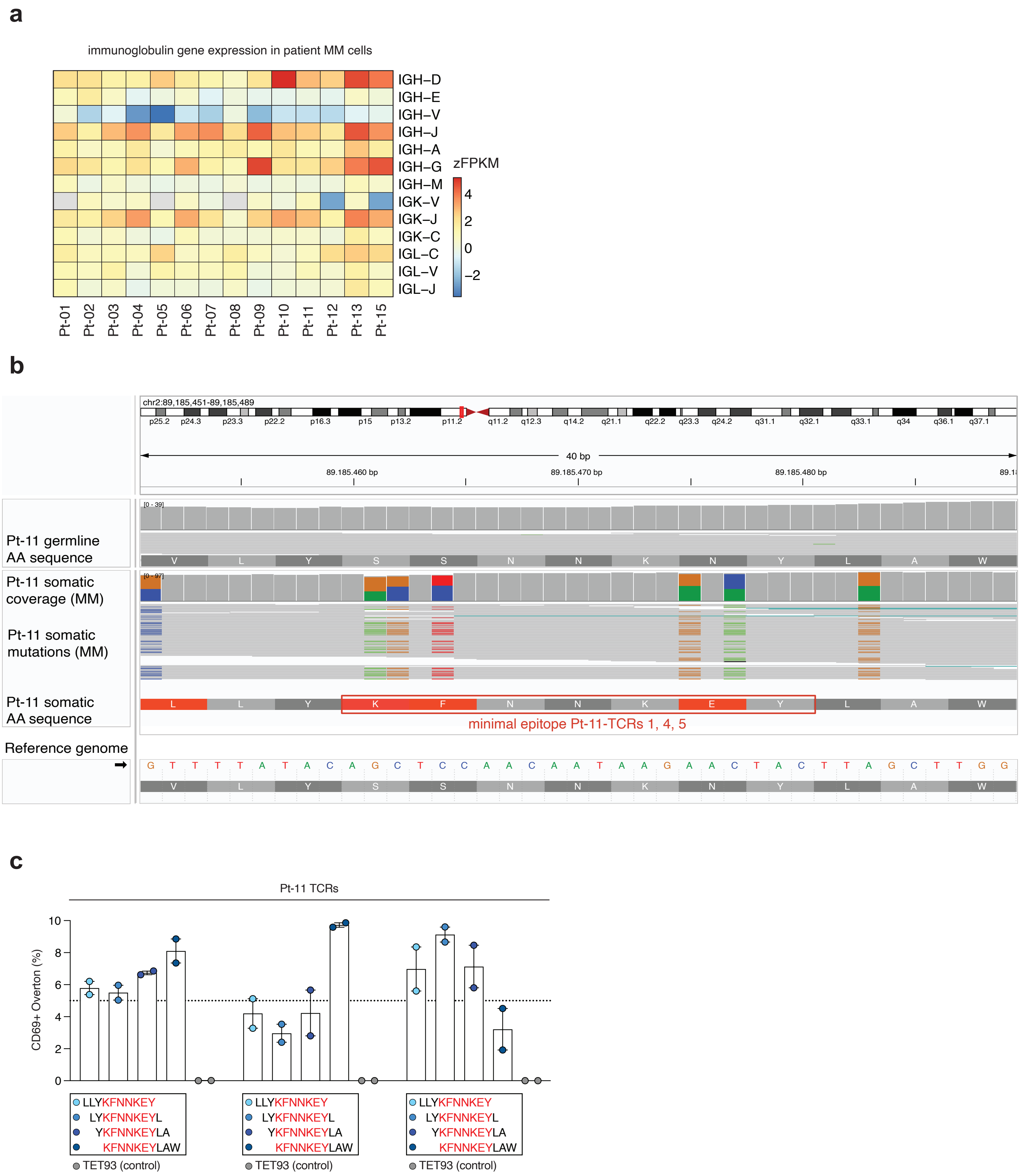
a
immunoglobulin gene expression in patient MM cells
IGH–D
IGH–E
IGH–V
IGH–J
IGH–A
IGH–G
IGH–M
IGK–V
IGK–J
IGK–C
IGL–C
IGL–V
IGL–J
zFPKM
4
2
0
−2
Pt-01
Pt-02
Pt-03
Pt-04
Pt-05
Pt-06
Pt-07
Pt-08
Pt-09
Pt-10
Pt-11
Pt-12
Pt-13
Pt-15
b
chr2:89,185,451-89,185,489
p25.2 p24.3 p23.3 p22.2 p16.3 p15 p13.2 p11.2 q11.2 q12.3 q14.2 q21.1 q22.2 q23.3 q24.2 q31.1 q32.1 q33.1 q34 q36.1 q37.1
40 bp
89.185.460 bp
89.185.470 bp
89.185.480 bp
[0 - 39]
Pt-11 germline
AA sequence
V L Y S S N N K N Y L A W
[0 - 97]
Pt-11 somatic
coverage (MM)
Pt-11 somatic
mutations (MM)
Pt-11 somatic
AA sequence
L L Y K F N N K E Y L A W
minimal epitope Pt-11-TCRs 1, 4, 5
Reference genome
G T T T T A T A C A G C T C C A A C A A T A A G A A C T A C T T A G C T T G G
V L Y S S N N K N Y L A W
c
Pt-11 TCRs
CD69+ Overton (%)
10
8
6
4
2
0
LLYKFNNKEY
LYKFNNKEYL
YKFNNKEYLA
KFNNKEYLAW
TET93 (control)

**Figure S7. Identification of a myeloma-specific neoepitope derived from immunoglobulin variable regions.**

**a,** Heatmap of z-score-normalized FPKM values of immunoglobulin gene expression in bulk RNA-seq from purified $CD138^{+}$ MM cells.
**b,** T-FINDER-based peptide scanning of four overlapping 10-mer peptides spanning the LLYKFNNKEYLAW sequence for reactivity by Pt-11-derived TCRs 1, 4, and 5.
**c,** DNA sequencing traces of the IGKV4-1 genomic locus in germline (PBMCs) and tumor (MM cells) DNA, showing somatic mutations and the location of the minimal reactive epitope in Pt-11-derived TCRs 1, 4, and 5.

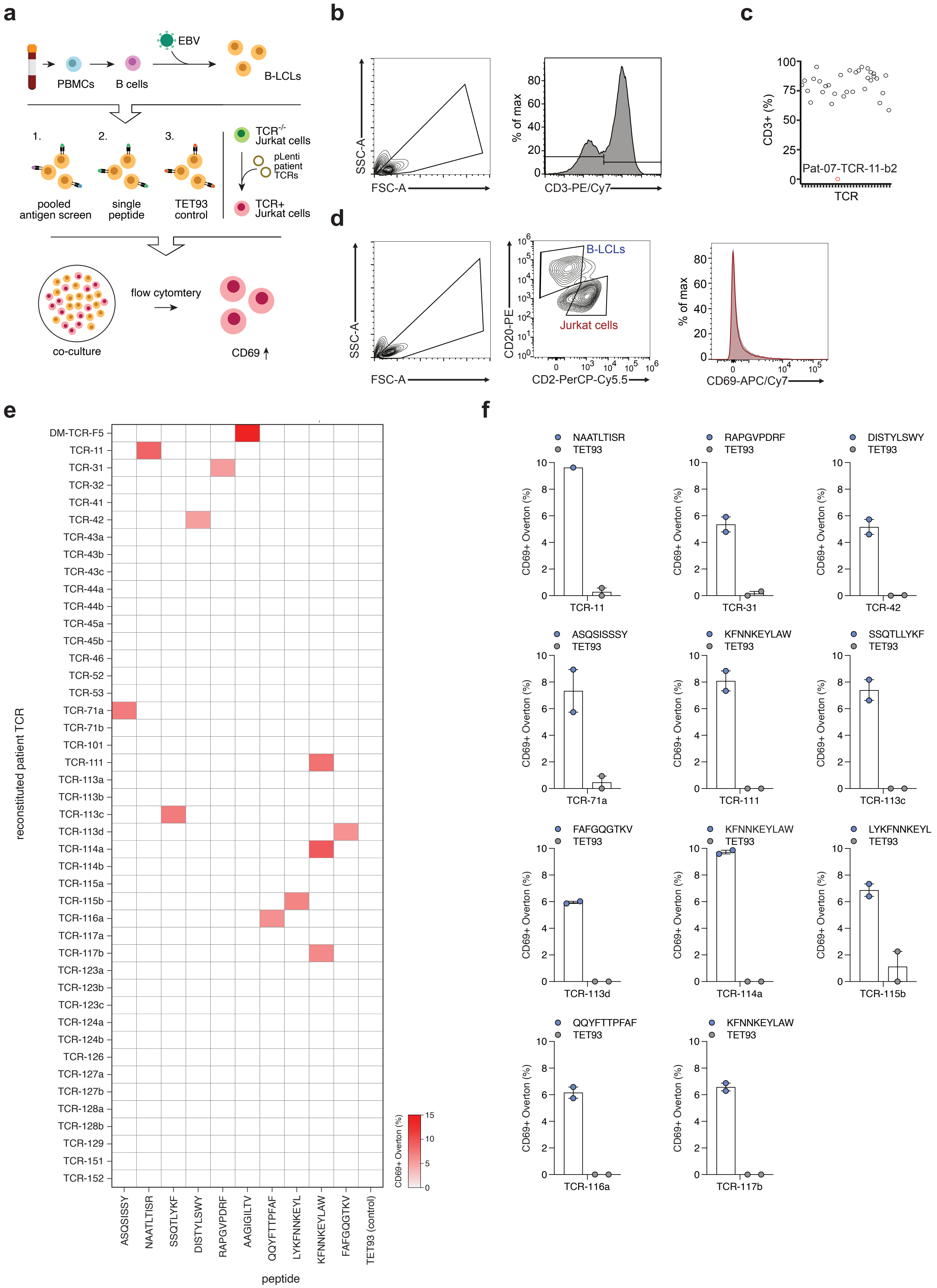

a
EBV
PBMCs
B cells
B-LCLs
1.
2.
3.
pooled antigen screen
single peptide
TET93 control
TCR-/- Jurkat cells
pLenti patient TCRs
TCR+ Jurkat cells
co-culture
flow cytomtery
CD69
b
SSC-A
FSC-A
% of max
CD3-PE/Cy7
c
CD3+ (%)
Pat-07-TCR-11-b2
TCR
d
B-LCLs
Jurkat cells
CD20-PE
CD2-PerCP-Cy5.5
CD69-APC/Cy7
e
reconstituted patient TCR
DM-TCR-F5
TCR-11
TCR-31
TCR-32
TCR-41
TCR-42
TCR-43a
TCR-43b
TCR-43c
TCR-44a
TCR-44b
TCR-45a
TCR-45b
TCR-46
TCR-52
TCR-53
TCR-71a
TCR-71b
TCR-101
TCR-111
TCR-113a
TCR-113b
TCR-113c
TCR-113d
TCR-114a
TCR-114b
TCR-115a
TCR-115b
TCR-116a
TCR-117a
TCR-117b
TCR-123a
TCR-123b
TCR-123c
TCR-124a
TCR-124b
TCR-126
TCR-127a
TCR-127b
TCR-128a
TCR-128b
TCR-129
TCR-151
TCR-152
ASQSISSY
NAATLTISR
SSQTLYKF
DISTYLSWY
RAPGVPDRF
AAGIGILTV
QQYFTTPFAF
LYKFNNKEYL
KFNNKEYLAW
FAFGQGTKV
TET93 (control)
peptide
CD69+ Overton (%)
f
NAATLTISR
TET93
TCR-11
RAPGVPDRF
TCR-31
DISTYLSWY
TCR-42
ASQSISSSY
TCR-71a
KFNNKEYLAW
TCR-111
SSQTLLYKF
TCR-113c
FAFGQGTKV
TCR-113d
TCR-114a
LYKFNNKEYL
TCR-115b
QQYFTTPFAF
TCR-116a
TCR-117b

**Figure S7. Antigen screening of tumor-reactive TCRs using the T-FINDER platform.**

**a,** Schematic of the T-FINDER assay: transgenic TCR-Jurkat reporter cells co-cultured with peptide-pulsed autologous B-LCLs. Activation measured by CD69 expression.
**b,** Flow cytometry gating strategy for detection of transduced Jurkat cells.
**c,** Dot plot of CD3 expression in transduced Jurkat cells confirming TCR surface expression (TCR 711c failed to express and was excluded).
**d,** Gating strategy for B-LCLs and reporter cells; CD69 analyzed using Overton subtraction relative to no peptide and TET93 irrelevant peptide controls.
**e,** Heatmap showing positive TCR:antigen pairs surpassing 5% CD69 Overton threshold relative to no peptide controls.
**f,** Quantification of TCR:antigen hits (N=2 co-cultures per combination). Corresponding irrelevant peptide control (TET93) included.

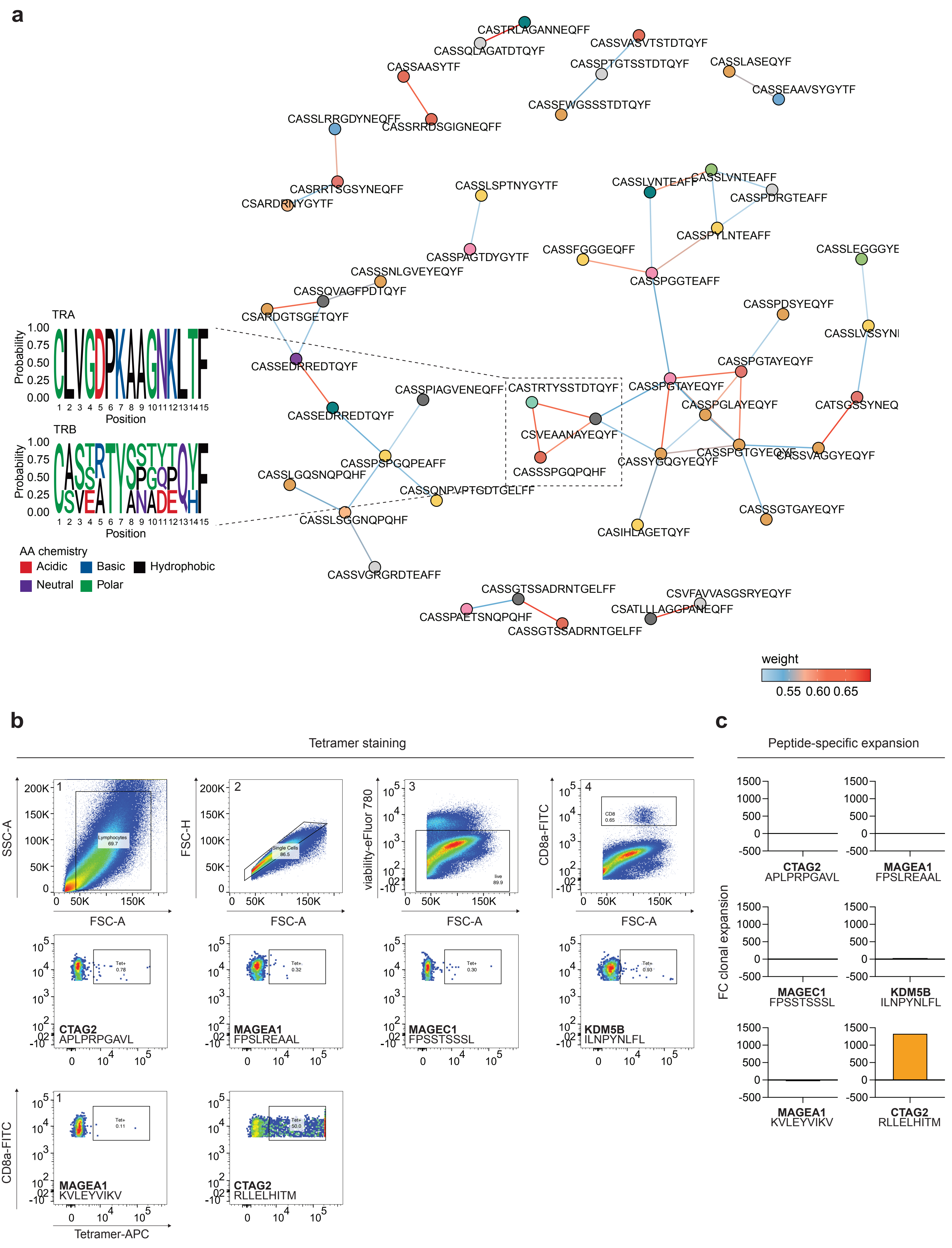
a
CASTRLAGANNEQFF
CASSQLAGATDTQYF
CASSVASVTSTDTQYF
CASSAASYTF
CASSPTGTSSTDTQYF
CASSLASEQYF
CASSEAAVSYGYTF
CASSFWGSSSTDTQYF
CASSLRRGDYNEQFF
CASSRRDSGIGNEQFF
CASRRTSGSYNEQFF
CSARDRNYGYTF
CASSLSPTNYGYTF
CASSLVNTEAFF
CASSLVNTEAFF
CASSPDRGTEAFF
CASSPYLNTEAFF
CASSPAGTDYGYTF
CASSFGGGEQFF
CASSLEGGGYE
CASSPGGTEAFF
CASSSNLGVEYEQYF
CASSQVAGFPDTQYF
CSARDGTSGETQYF
CASSPDSYEQYF
CASSLVSSYNI
TRA
Probability
1.00
0.75
0.50
0.25
0.00
Position
CASSEDRREDTQYF
CASSPGTAYEQYF
CASSPIAGVENEQFF
CASTRTYSSTDTQYF
CASSPGTAYEQYF
CASSPGLAYEQYF
CATSGSSYNEQ
CASSEDRREDTQYF
CSVEAANAYEQYF
TRB
CASSPGTGYEQYF
CASSVAGGYEQYF
CASSYGQGYEQYF
CASSPSPGQPEAFF
CASSSPGQPQHF
CASSLGQSNQPQHF
CASSQNPVPTGDTGELFF
CASSSGTGAYEQYF
CASSLSGGNQPQHF
CASIHLAGETQYF
AA chemistry
Acidic
Basic
Hydrophobic
Neutral
Polar
CASSVGRGRDTEAFF
CASSGTSSADRNTGELFF
CSVFAVVASGSRYEQYF
CSATLLLAGGPANEQFF
CASSPAETSNQPQHF
CASSGTSSADRNTGELFF
weight
0.55 0.60 0.65
b
Tetramer staining
SSC-A
FSC-A
Lymphocytes
69.7
FSC-H
Single Cells
86.5
viability-eFluor 780
live
89.9
CD8a-FITC
CD8
0.65
Tet+
0.78
CTAG2
APLPRPGAVL
Tet+
0.32
MAGEA1
FPSLREAAL
Tet+
0.30
MAGEC1
FPSSTSSSL
Tet+
0.93
KDM5B
ILNPYNLFL
CD8a-FITC
Tet+
0.11
MAGEA1
KVLEYVIKV
Tet+
50.0
CTAG2
RLLELHITM
Tetramer-APC
c
Peptide-specific expansion
FC clonal expansion
1500
1000
500
0
-500
CTAG2
APLPRPGAVL
MAGEA1
FPSLREAAL
MAGEC1
FPSSTSSSL
KDM5B
ILNPYNLFL
MAGEA1
KVLEYVIKV
CTAG2
RLLELHITM

**Figure S9. Tumor-reactive TCR convergence across patients.**

**a,** Network of TCRs with similar CDR3αβ sequences based on pairwise BLOSUM45 similarity. Edges shown only above 95% bootstrap threshold. Nodes colored by patient of origin (N=15).
**b,** Flow cytometry plots of peptide-MHC tetramer staining for immunopeptidomics-derived shared peptides in three patients with convergent TCRs.
**c,** Fold change in clonotype frequency from day 0 to day 28 in the MANAFEST assay, where peptide-loaded autologous PBMCs served as APCs.

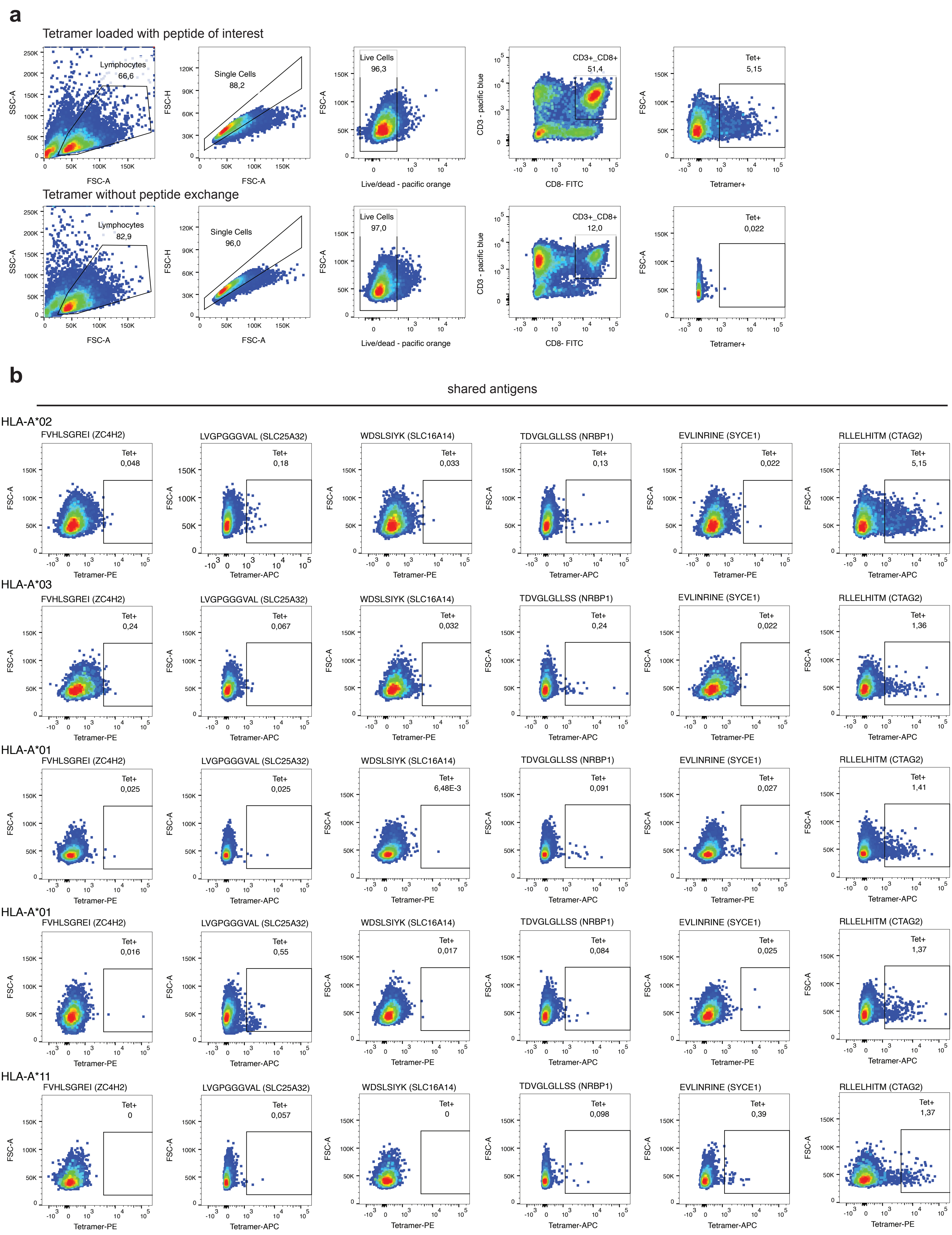

a
Tetramer loaded with peptide of interest
Lymphocytes
66,6
SSC-A
FSC-A
Single Cells
88,2
FSC-H
FSC-A
Live Cells
96,3
FSC-A
Live/dead - pacific orange
CD3+_CD8+
51,4
CD3 - pacific blue
CD8- FITC
Tet+
5,15
FSC-A
Tetramer+
Tetramer without peptide exchange
Lymphocytes
82,9
SSC-A
FSC-A
Single Cells
96,0
FSC-H
FSC-A
Live Cells
97,0
FSC-A
Live/dead - pacific orange
CD3+_CD8+
12,0
CD3 - pacific blue
CD8- FITC
Tet+
0,022
FSC-A
Tetramer+
b
shared antigens
HLA-A*02
FVHLSGREI (ZC4H2)
Tet+
0,048
Tetramer-PE
LVGPGGGVAL (SLC25A32)
Tet+
0,18
Tetramer-APC
WDSLSIYK (SLC16A14)
Tet+
0,033
Tetramer-PE
TDVGLGLLSS (NRBP1)
Tet+
0,13
Tetramer-APC
EVLINRINE (SYCE1)
Tet+
0,022
Tetramer-PE
RLLELHITM (CTAG2)
Tet+
5,15
Tetramer-APC
HLA-A*03
FVHLSGREI (ZC4H2)
Tet+
0,24
Tetramer-PE
LVGPGGGVAL (SLC25A32)
Tet+
0,067
Tetramer-APC
WDSLSIYK (SLC16A14)
Tet+
0,032
Tetramer-PE
TDVGLGLLSS (NRBP1)
Tet+
0,24
Tetramer-APC
EVLINRINE (SYCE1)
Tet+
0,022
Tetramer-PE
RLLELHITM (CTAG2)
Tet+
1,36
Tetramer-APC
HLA-A*01
FVHLSGREI (ZC4H2)
Tet+
0,025
Tetramer-PE
LVGPGGGVAL (SLC25A32)
Tet+
0,025
Tetramer-APC
WDSLSIYK (SLC16A14)
Tet+
6,48E-3
Tetramer-PE
TDVGLGLLSS (NRBP1)
Tet+
0,091
Tetramer-APC
EVLINRINE (SYCE1)
Tet+
0,027
Tetramer-PE
RLLELHITM (CTAG2)
Tet+
1,41
Tetramer-APC
HLA-A*01
FVHLSGREI (ZC4H2)
Tet+
0,016
Tetramer-PE
LVGPGGGVAL (SLC25A32)
Tet+
0,55
Tetramer-APC
WDSLSIYK (SLC16A14)
Tet+
0,017
Tetramer-PE
TDVGLGLLSS (NRBP1)
Tet+
0,084
Tetramer-APC
EVLINRINE (SYCE1)
Tet+
0,025
Tetramer-PE
RLLELHITM (CTAG2)
Tet+
1,37
Tetramer-APC
HLA-A*11
FVHLSGREI (ZC4H2)
Tet+
0
Tetramer-PE
LVGPGGGVAL (SLC25A32)
Tet+
0,057
Tetramer-APC
WDSLSIYK (SLC16A14)
Tet+
0
Tetramer-PE
TDVGLGLLSS (NRBP1)
Tet+
0,098
Tetramer-APC
EVLINRINE (SYCE1)
Tet+
0,39
Tetramer-APC
RLLELHITM (CTAG2)
Tet+
1,37
Tetramer-PE
FSC-A

**Figure S10. HLA binding promiscuity of shared immunopeptidome-derived antigens.**

**a,** Gating strategy for peptide-MHC tetramer staining of autologous BMLs.
**b,** Representative tetramer staining for shared peptides presented by different HLA-A supergroups (HLA-A*01, *02, *03, *11) in PBMCs from 5 healthy donors.

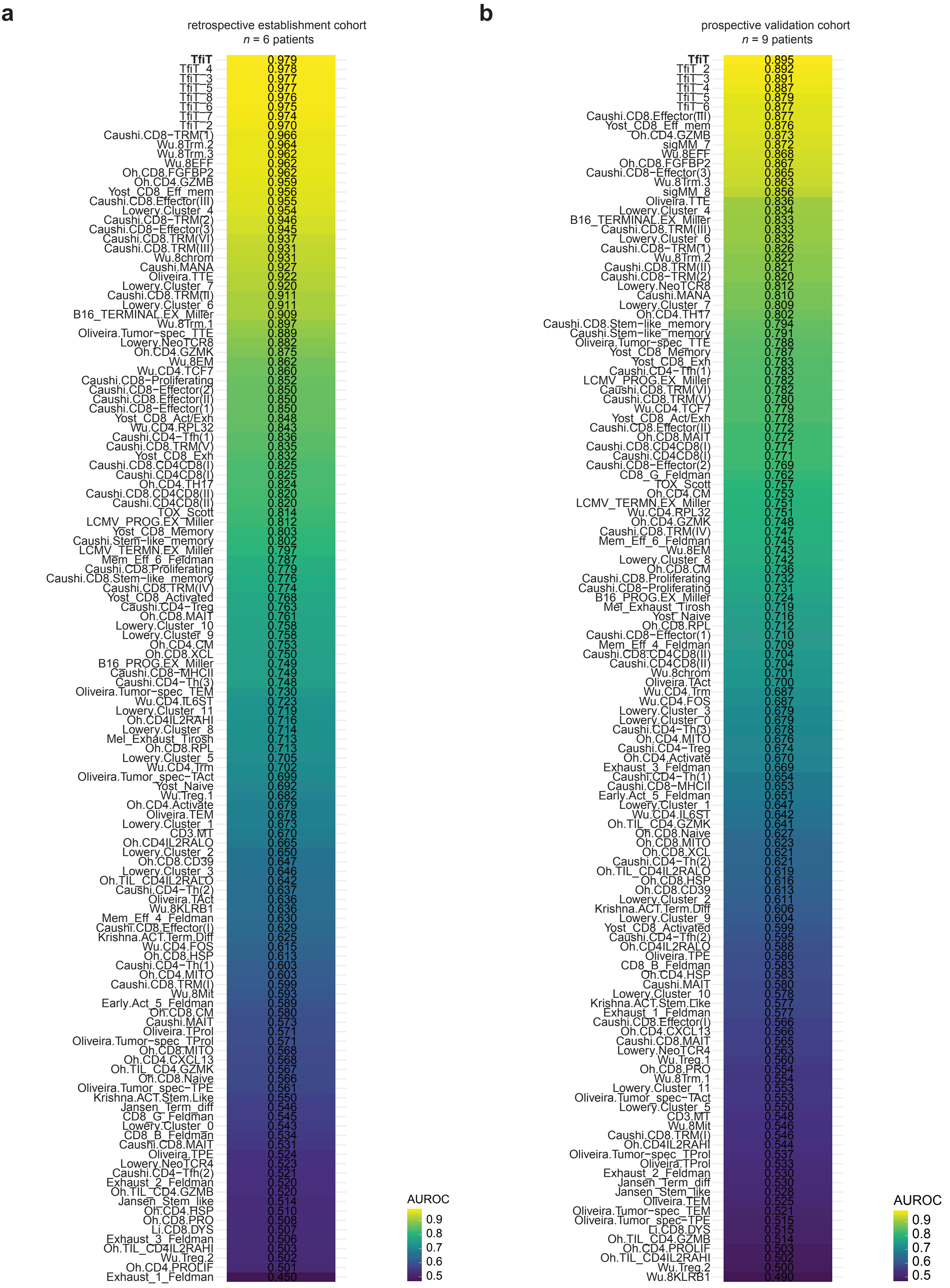
a
retrospective establishment cohort
n = 6 patients
TfiT 0.979
TfiT_4 0.978
TfiT_3 0.977
TfiT_5 0.977
TfiT_8 0.976
TfiT_6 0.975
TfiT_7 0.974
TfiT_2 0.970
Caushi.CD8−TRM(1) 0.966
Wu.8Trm.2 0.964
Wu.8Trm.3 0.962
Wu.8EFF 0.962
Oh.CD8.FGFBP2 0.962
Oh.CD4.GZMB 0.959
Yost_CD8_Eff_mem 0.956
Caushi.CD8.Effector(III) 0.955
Lowery.Cluster_4 0.954
Caushi.CD8−TRM(2) 0.946
Caushi.CD8−Effector(3) 0.945
Caushi.CD8.TRM(VI) 0.937
Caushi.CD8.TRM(III) 0.931
Wu.8chrom 0.931
Caushi.MANA 0.927
Oliveira.TTE 0.922
Lowery.Cluster_7 0.920
Caushi.CD8.TRM(II) 0.911
Lowery.Cluster_6 0.911
B16_TERMINAL.EX_Miller 0.909
Wu.8Trm.1 0.897
Oliveira.Tumor−spec_TTE 0.889
Lowery.NeoTCR8 0.882
Oh.CD4.GZMK 0.875
Wu.8EM 0.862
Wu.CD4.TCF7 0.860
Caushi.CD8−Proliferating 0.852
Caushi.CD8−Effector(2) 0.850
Caushi.CD8.Effector(II) 0.850
Caushi.CD8−Effector(1) 0.850
Yost_CD8_Act/Exh 0.848
Wu.CD4.RPL32 0.843
Caushi.CD4−Tfh(1) 0.836
Caushi.CD8.TRM(V) 0.835
Yost_CD8_Exh 0.832
Caushi.CD8.CD4CD8(I) 0.825
Caushi.CD4CD8(I) 0.825
Oh.CD4.TH17 0.824
Caushi.CD8.CD4CD8(II) 0.820
Caushi.CD4CD8(II) 0.820
TOX_Scott 0.814
LCMV_PROG.EX_Miller 0.812
Yost_CD8_Memory 0.803
Caushi.Stem−like_memory 0.802
LCMV_TERMN.EX_Miller 0.797
Mem_Eff_6_Feldman 0.787
Caushi.CD8.Proliferating 0.779
Caushi.CD8.Stem−like_memory 0.776
Caushi.CD8.TRM(IV) 0.774
Yost_CD8_Activated 0.768
Caushi.CD4−Treg 0.763
Oh.CD8.MAIT 0.761
Lowery.Cluster_10 0.758
Lowery.Cluster_9 0.758
Oh.CD4.CM 0.753
Oh.CD8.XCL 0.750
B16_PROG.EX_Miller 0.749
Caushi.CD8−MHCII 0.749
Caushi.CD4−Th(3) 0.748
Oliveira.Tumor−spec_TEM 0.730
Wu.CD4.IL6ST 0.723
Lowery.Cluster_11 0.719
Oh.CD4IL2RAHI 0.716
Lowery.Cluster_8 0.714
Mel_Exhaust_Tirosh 0.713
Oh.CD8.RPL 0.713
Lowery.Cluster_5 0.705
Wu.CD4.Trm 0.702
Oliveira.Tumor_spec−TAct 0.699
Yost_Naive 0.692
Wu.Treg.1 0.682
Oh.CD4.Activate 0.679
Oliveira.TEM 0.678
Lowery.Cluster_1 0.673
CD3.MT 0.670
Oh.CD4IL2RALO 0.665
Lowery.Cluster_2 0.650
Oh.CD8.CD39 0.647
Lowery.Cluster_3 0.646
Oh.TIL_CD4IL2RALO 0.642
Caushi.CD4−Th(2) 0.637
Oliveira.TAct 0.636
Wu.8KLRB1 0.636
Mem_Eff_4_Feldman 0.630
Caushi.CD8.Effector(I) 0.629
Krishna.ACT.Term.Diff 0.625
Wu.CD4.FOS 0.615
Oh.CD8.HSP 0.613
Caushi.CD4−Th(1) 0.603
Oh.CD4.MITO 0.603
Caushi.CD8.TRM(I) 0.599
Wu.8Mit 0.593
Early.Act_5_Feldman 0.589
Oh.CD8.CM 0.580
Caushi.MAIT 0.573
Oliveira.TProl 0.571
Oliveira.Tumor−spec_TProl 0.571
Oh.CD8.MITO 0.568
Oh.CD4.CXCL13 0.568
Oh.TIL_CD4.GZMK 0.567
Oh.CD8.Naive 0.566
Oliveira.Tumor_spec−TPE 0.561
Krishna.ACT.Stem.Like 0.550
Jansen_Term_diff 0.546
CD8_G_Feldman 0.545
Lowery.Cluster_0 0.543
CD8_B_Feldman 0.534
Caushi.CD8.MAIT 0.531
Oliveira.TPE 0.524
Lowery.NeoTCR4 0.523
Caushi.CD4−Tfh(2) 0.521
Exhaust_2_Feldman 0.520
Oh.TIL_CD4.GZMB 0.520
Jansen_Stem_like 0.514
Oh.CD4.HSP 0.510
Oh.CD8.PRO 0.508
Li.CD8.DYS 0.507
Exhaust_3_Feldman 0.506
Oh.TIL_CD4IL2RAHI 0.503
Wu.Treg.2 0.502
Oh.CD4.PROLIF 0.501
Exhaust_1_Feldman 0.450
AUROC
0.9
0.8
0.7
0.6
0.5
b
prospective validation cohort
n = 9 patients
TfiT 0.895
TfiT_2 0.892
TfiT_3 0.891
TfiT_4 0.887
TfiT_5 0.879
TfiT_6 0.877
Caushi.CD8.Effector(III) 0.877
Yost_CD8_Eff_mem 0.876
Oh.CD4.GZMB 0.873
sigMM_7 0.872
Wu.8EFF 0.868
Oh.CD8.FGFBP2 0.867
Caushi.CD8−Effector(3) 0.865
Wu.8Trm.3 0.863
sigMM_8 0.856
Oliveira.TTE 0.836
Lowery.Cluster_4 0.834
B16_TERMINAL.EX_Miller 0.833
Caushi.CD8.TRM(III) 0.833
Lowery.Cluster_6 0.832
Caushi.CD8−TRM(1) 0.826
Wu.8Trm.2 0.822
Caushi.CD8.TRM(II) 0.821
Caushi.CD8−TRM(2) 0.820
Lowery.NeoTCR8 0.812
Caushi.MANA 0.810
Lowery.Cluster_7 0.809
Oh.CD4.TH17 0.802
Caushi.CD8.Stem−like_memory 0.794
Caushi.Stem−like_memory 0.791
Oliveira.Tumor−spec_TTE 0.788
Yost_CD8_Memory 0.787
Yost_CD8_Exh 0.783
Caushi.CD4−Tfh(1) 0.783
LCMV_PROG.EX_Miller 0.782
Caushi.CD8.TRM(VI) 0.782
Caushi.CD8.TRM(V) 0.780
Wu.CD4.TCF7 0.779
Yost_CD8_Act/Exh 0.778
Caushi.CD8.Effector(II) 0.772
Oh.CD8.MAIT 0.772
Caushi.CD8.CD4CD8(I) 0.771
Caushi.CD4CD8(I) 0.771
Caushi.CD8−Effector(2) 0.769
CD8_G_Feldman 0.762
TOX_Scott 0.757
Oh.CD4.CM 0.753
LCMV_TERMN.EX_Miller 0.751
Wu.CD4.RPL32 0.751
Oh.CD4.GZMK 0.748
Caushi.CD8.TRM(IV) 0.747
Mem_Eff_6_Feldman 0.745
Wu.8EM 0.743
Lowery.Cluster_8 0.742
Oh.CD8.CM 0.736
Caushi.CD8.Proliferating 0.732
Caushi.CD8−Proliferating 0.731
B16_PROG.EX_Miller 0.724
Mel_Exhaust_Tirosh 0.719
Yost_Naive 0.716
Oh.CD8.RPL 0.712
Caushi.CD8−Effector(1) 0.710
Mem_Eff_4_Feldman 0.709
Caushi.CD8.CD4CD8(II) 0.704
Caushi.CD4CD8(II) 0.704
Wu.8chrom 0.701
Oliveira.TAct 0.700
Wu.CD4.Trm 0.687
Wu.CD4.FOS 0.687
Lowery.Cluster_3 0.679
Lowery.Cluster_0 0.679
Caushi.CD4−Th(3) 0.678
Oh.CD4.MITO 0.676
Caushi.CD4−Treg 0.674
Oh.CD4.Activate 0.670
Exhaust_3_Feldman 0.669
Caushi.CD4−Th(1) 0.654
Caushi.CD8−MHCII 0.653
Early.Act_5_Feldman 0.651
Lowery.Cluster_1 0.647
Wu.CD4.IL6ST 0.642
Oh.TIL_CD4.GZMK 0.641
Oh.CD8.Naive 0.627
Oh.CD8.MITO 0.623
Oh.CD8.XCL 0.621
Caushi.CD4−Th(2) 0.621
Oh.TIL_CD4IL2RALO 0.619
Oh.CD8.HSP 0.616
Oh.CD8.CD39 0.613
Lowery.Cluster_2 0.611
Krishna.ACT.Term.Diff 0.606
Lowery.Cluster_9 0.604
Yost_CD8_Activated 0.599
Caushi.CD4−Tfh(2) 0.595
Oh.CD4IL2RALO 0.588
Oliveira.TPE 0.586
CD8_B_Feldman 0.583
Oh.CD4.HSP 0.583
Caushi.MAIT 0.580
Lowery.Cluster_10 0.578
Krishna.ACT.Stem.Like 0.577
Exhaust_1_Feldman 0.577
Caushi.CD8.Effector(I) 0.566
Oh.CD4.CXCL13 0.566
Caushi.CD8.MAIT 0.565
Lowery.NeoTCR4 0.563
Wu.Treg.1 0.560
Oh.CD8.PRO 0.554
Wu.8Trm.1 0.554
Lowery.Cluster_11 0.553
Oliveira.Tumor_spec−TAct 0.553
Lowery.Cluster_5 0.550
CD3.MT 0.548
Wu.8Mit 0.546
Caushi.CD8.TRM(I) 0.546
Oh.CD4IL2RAHI 0.544
Oliveira.Tumor−spec_TProl 0.537
Oliveira.TProl 0.533
Exhaust_2_Feldman 0.530
Jansen_Term_diff 0.530
Jansen_Stem_like 0.528
Oliveira.TEM 0.525
Oliveira.Tumor−spec_TEM 0.521
Oliveira.Tumor_spec−TPE 0.515
Li.CD8.DYS 0.515
Oh.TIL_CD4.GZMB 0.514
Oh.CD4.PROLIF 0.503
Oh.TIL_CD4IL2RAHI 0.502
Wu.Treg.2 0.500
Wu.8KLRB1 0.490
AUROC
0.9
0.8
0.7
0.6
0.5

**Figure S11. Benchmarking of TfiT transcriptional signature for tumor-reactive TCR prediction.**

**a,** Heatmap showing AUROC values of 122 published and 8 newly generated T cell transcriptional signatures in predicting tumor-reactive clonotypes in BMLs from 6 NDMM patients (retrospective cohort). **b,** As in (a), evaluated in BMLs from 9 NDMM patients in the prospective validation cohort.

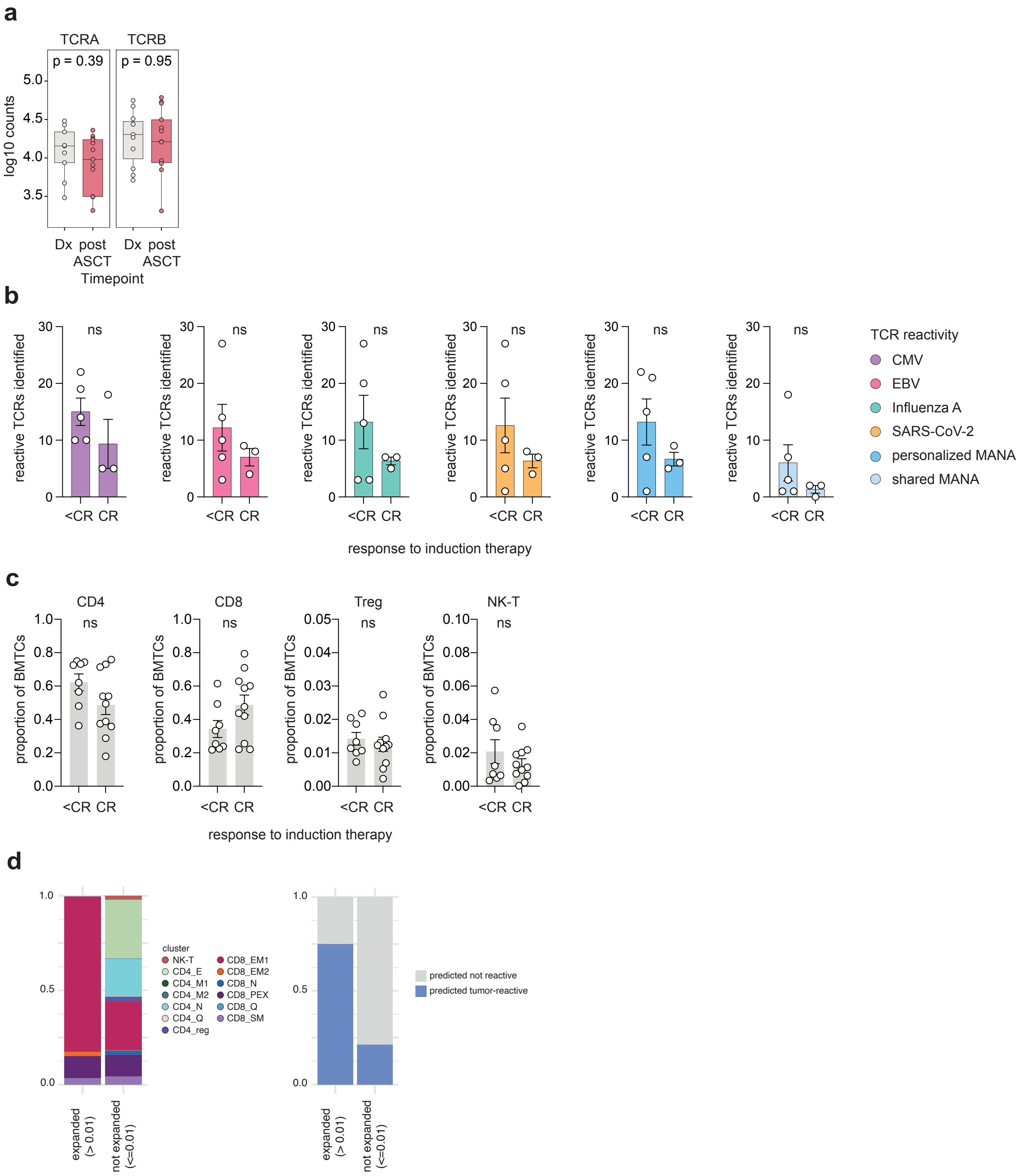
a
TCRA
TCRB
p = 0.39
p = 0.95
log10 counts
5.0
4.5
4.0
3.5
Dx post ASCT
Dx post ASCT
Timepoint
b
reactive TCRs identified
ns
<CR CR
response to induction therapy
TCR reactivity
CMV
EBV
Influenza A
SARS-CoV-2
personalized MANA
shared MANA
c
CD4
CD8
Treg
NK-T
ns
proportion of BMTCs
<CR CR
response to induction therapy
d
cluster
NK-T
CD4_E
CD4_M1
CD4_M2
CD4_N
CD4_Q
CD4_reg
CD8_EM1
CD8_EM2
CD8_N
CD8_PEX
CD8_Q
CD8_SM
expanded (> 0.01)
not expanded (<=0.01)
predicted not reactive
predicted tumor-reactive

**Figure S12. Clinical associations of BML T cell features in newly diagnosed multiple myeloma.**

**a,** Log10 transformed counts of TCR alpha and beta transcript counts before and post therapy in longitudinal CARDAMON-cohort[78].
**b,** Number of tumor-reactive, virus-reactive, and orphan TCRs as determined by MANAFEST Assay analysis per patient at diagnosis stratified by response to induction therapy (complete response [CR] vs. non-CR). Statistical significance was assessed using unpaired two-tailed t-test with Welch's correction.
**c,** Frequency of selected T cell subsets in BMLs at baseline, stratified by remission status following induction therapy. Statistical analysis as in (a).
**d,** BML cluster distribution (left) and predicted tumor-reactivity distribution (right) in bsAb cohort68 prior to therapy by binary clonotype expansion category (proportion > or < 0.01).
